# YAP/TAZ enhance P-body formation to promote tumorigenesis

**DOI:** 10.1101/2023.06.02.542626

**Authors:** Xia Shen, Xiang Peng, Yuegui Guo, Zhujiang Dai, Long Cui, Wei Yu, Yun Liu, Chen-Ying Liu

**Author notes:** Correspondence and requests for materials should be addressed to Yun Liu or Chen-Ying Liu. These authors contribute equally (Xia Shen, Xiang Peng, Yuegui Guo).

## Abstract

The role of Processing bodies (P-bodies) in tumorigenesis and tumor progression is not well understood. Here, we showed that the oncogenes YAP/TAZ promote P-body formation in a series of cancer cell lines. Mechanistically, both transcriptional activation of the P-body-related genes SAMD4A, AJUBA, and WTIP and transcriptional suppression of the tumor suppressor gene PNRC1 are involved in enhancing the effects of YAP/TAZ on P-body formation in CRC cells. By reexpression of PNRC1 or knockdown of P-body core genes (DDX6, DCP1A, and LSM14A), we determined that disruption of P-bodies attenuates cell proliferation, cell migration and tumor growth induced by overexpression of YAP^5SA^ in CRC. Analysis of a pancancer CRISPR screen database (DepMap) revealed codependencies between YAP/TEAD and the P-body core genes and correlations between the mRNA levels of SAMD4A, AJUBA, WTIP, PNRC1 and YAP target genes. Our study suggests that the P-body is a new downstream effector of YAP/TAZ, which implies that reexpression of PNRC1 or disruption of P-bodies is a potential therapeutic strategy for tumors with active YAP.

## Introduction

The Hippo pathway is an evolutionally conserved signaling pathway that regulates organ size and plays vital roles in development and tissue homeostasis (Driskill & Pan, 2021; Ma et al, 2019; Russell & Camargo, 2022). The transcriptional output of the Hippo pathway is mainly mediated by the YAP/TAZ-TEAD transcription complex. In response to various extracellular or intracellular signals, including cell–cell contact, mechanical force, serum stimulation, cellular stress and cellular energy status, the YAP/TAZ-TEAD complex modulates target gene expression to respond to environmental cues(Calvo et al, 2013; Misra & Irvine, 2018; Yu et al, 2012; Zhao et al, 2007). Although initially identified as transcriptional coactivators, YAP/TAZ can also function as corepressors to inhibit target gene transcription by recruiting the nucleosome remodeling and histone deacetylase (NuRD) complex (Kim et al, 2015b). The evidence of Hippo pathway dysregulation in a variety of cancers and the list of YAP/TAZ target genes continue to increase (Calses et al, 2019; Kulkarni et al, 2020; Nguyen & Yi, 2019; Wang et al, 2018; Zanconato et al, 2016). Dysregulation of YAP/TAZ-TEAD transcriptional output endows tumor cells with every hallmark of cancer, including sustained proliferation, resistance to apoptosis, tumor-promoting inflammation, tumor immune escape, dysregulated tumor metabolism, etc. (Calses et al, 2019; Hanahan, 2022; Kulkarni et al, 2020; Nguyen & Yi, 2019; Zanconato et al, 2016).

At the cellular organization level, the YAP/TAZ-TEAD transcription complex modulates mitochondrial fusion; cytoskeleton, primary cilium, and focal adhesion assembly; and caveolae formation (Kim et al, 2015a; Mason et al, 2019; Nagaraj et al, 2012; Qiao et al, 2017; Rausch et al, 2019). Processing bodies (P-bodies) are cytoplasmic membraneless organelles that consist of ribonucleoprotein complexes (RNPs) and are formed by phase separation (Luo et al, 2018; Riggs et al, 2020). Although initial studies hypothesized that mRNAs in P-bodies are targeted for decay and translational repression, it was subsequently suggested that P-bodies are not required for mRNA decay and that repressed mRNAs can be recycled from P-bodies to reenter translation; thus, the primary function of P-bodies is controlling the storage of untranslated mRNAs (Decker & Parker, 2012; Hubstenberger et al, 2017; Luo et al, 2018). The role of P-bodies in tumorigenesis and tumor progression is not well studied (Anderson et al, 2015; Lavalee et al, 2021; Riggs et al, 2020). The formation of P-bodies is correlated with epithelial-mesenchymal transition (EMT) in breast cancer (Hardy et al, 2017). In contrast, there is also evidence that an increase in P-bodies leads to attenuated growth, migration and invasion of prostate cancer cells (Bearss et al, 2021). Recently, YAP was reported to be a negative regulator of P-bodies and to be involved in Kaposi sarcoma-associated herpesvirus (KHSV)-induced P-body disassembly in human umbilical vein endothelial cells (HUVECs) (Castle et al, 2021). However, this regulatory axis and the potential function of P-bodies in YAP-induced tumorigenesis remain unclear.

In this study, we discovered that YAP/TAZ are enhancers but not negative regulators of P-body formation in a series of cancer cell lines. YAP/TAZ modulate the transcription of multiple P-body-related genes, especially repressing the transcription of the tumor suppressor proline-rich nuclear receptor coactivator 1 (PNRC1) through cooperation with the NuRD complex. As a direct YAP/TAZ target gene, PNRC1 functions as a critical effector in YAP-induced biogenesis of P-bodies and tumorigenesis in colorectal cancer (CRC). Furthermore, disruption of P-bodies by knockdown of core component genes of P-bodies attenuated the protumorigenic effects of YAP in CRC. Thus, our study reveals a YAP–P-body positive regulatory axis in colorectal cancer, which exposes the vital role of YAP/TAZ in the biogenesis of P-bodies in tumors and implies that reexpression of PNRC1 or disruption of P-bodies is a potential therapeutic strategy for cancers with active YAP.

## Results

### YAP/TAZ regulates the transcription of P-body-related genes

Previously, to identify the new target genes and molecular signatures of YAP/TAZ in colorectal cancer, we performed RNA sequencing analysis of HCT116 CRC cells with simultaneous knockdown of YAP and TAZ (GSE176475) (Guo et al, 2022). Gene ontology enrichment analysis of the 674 differentially expressed genes upon knockdown of YAP/TAZ (FC>2, p<0.05) revealed that the genes downregulated by YAP/TAZ knockdown were enriched in the term P-body in the cellular component category (Figure 1A). We further expanded our analysis to the moderately differentially expressed genes (FC<0.66 or >1.5) that were annotated as related to P-bodies (Figure 1-Figure Supplement 1A, Appendix Table S1). Through integration with the public ChIP-seq data for TEAD4 in HCT116 cells from the ENCODE database, AJUBA, WTIP, NOCT, SAMD4A and PNRC1 were selected for in-depth investigation (Figure 1B, Figure 1-Figure Supplement 1A). Intriguingly, the public TEAD4 ChIP-seq datasets for the other three cancer cell lines (A549, MCF7, and MDA-MB-231), not just HCT116 cells, also showed strong TEAD4 binding peaks in the genomic loci of these five P-body-related genes (Figure 1-Figure Supplement 1B) (Mei et al, 2017). HCT116, A549, MCF7, and MDA-MB-231 are well established cell models for exploring YAP/TAZ function and the cell proliferation of these four cell lines are dependent on YAP/TAZ activity (Rosenbluh et al, 2012; Shreberk-Shaked et al, 2020; Zanconato et al, 2015; Zhu et al, 2019). It is worth noting cell contact inhibition were observed in HCT116 and MDA-MB-231 and YAP remains in the nucleus regardless of cell–cell contact in A549 and MCF7 cells (Fan et al, 2013; Kim et al, 2011; Lee et al, 2018; Wu et al, 2019). The ChIP–qPCR results in HCT116 cells further confirmed that TEAD4 bound to the promoter regions of AJUBA, WTIP, NOCT, SAMD4A and PNRC1 and to the intronic region of PNRC1 (Figure 1B). Next, we confirmed the significantly downregulated mRNA expression of AJUBA, WTIP, SAMD4A and NOCT and moderately increased expression of PNRC1 in YAP/TAZ knockdown HCT116 cells by qPCR analysis; this pattern was also observed in A549 lung cancer cells and MDA-MB-231 breast cancer cells (Figure 1C). Consistent with these findings, overexpression of the constitutively active YAP^5SA^ mutant but not the TEAD binding-defective YAP^5SA-S94A^ mutant significantly decreased the mRNA level of PNRC1 and increased the mRNA level of SAMD4A in HCT116, MCF7 and A549 cells (Figure 1D). Enhanced expression of AJUBA and WTIP was observed in HCT116 and MCF7 cells but not in A549 cells (Figure 1D). Since NOCT was not affected by overexpression of YAP^5SA^ in either MCF7 or A549 cells, we did not investigate NOCT in subsequent functional experiments (Figure 1D). Finally, we confirmed that the protein level of PNRC1 was increased by knockdown of YAP/TAZ in HCT116 cells (Figure 1-Figure Supplement 1C). Additionally, overexpression of YAP^5SA^ but not YAP^5SA-S94A^ decreased the protein level of PNRC1 in HCT116, A549 and MDA-MB-231 cells (Figure 1-Figure Supplement 1D). Overall, these data demonstrate that YAP/TAZ modulate the transcription of P-body-related genes through the TEAD transcription factors.

**Figure 1.**
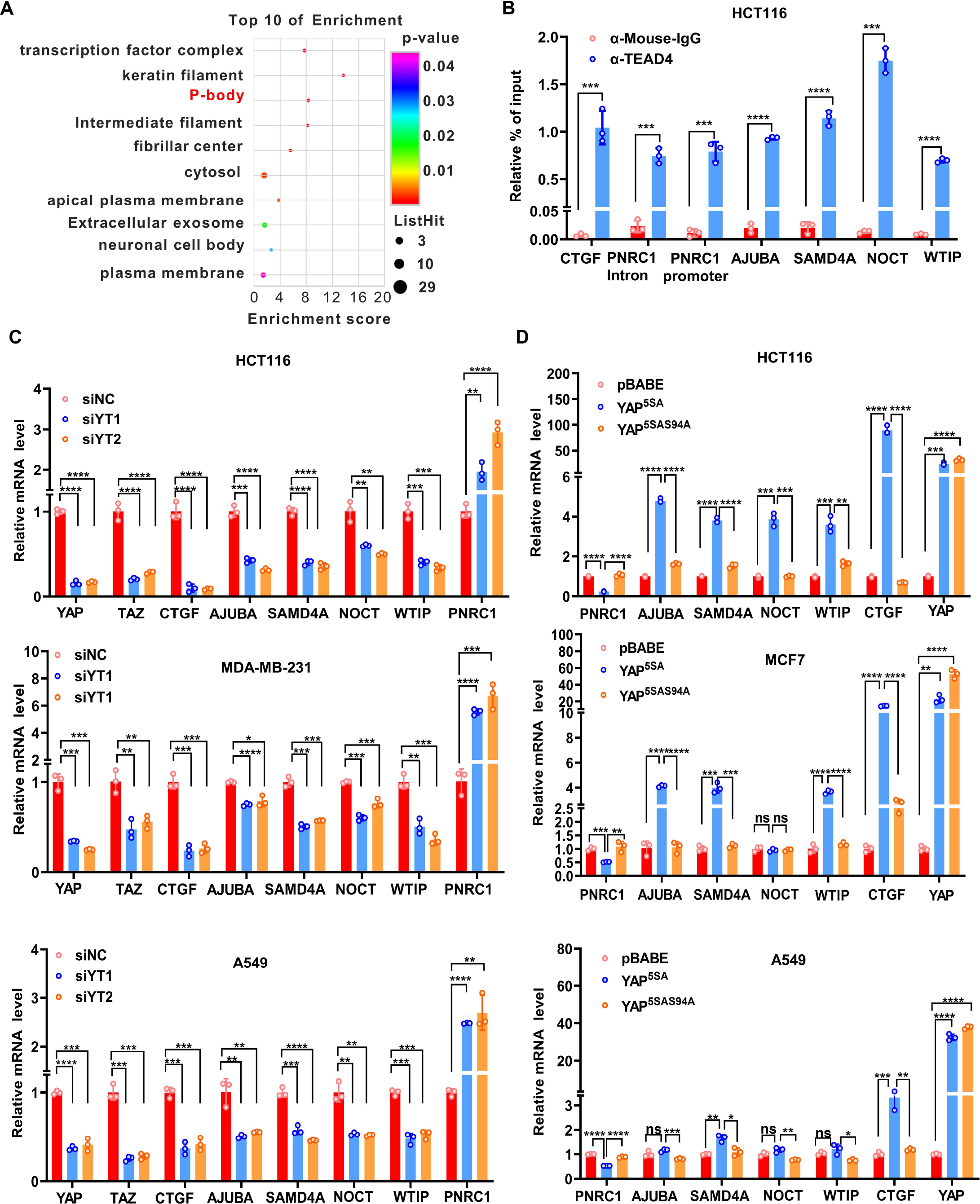
YAP/TAZ transcriptionally regulates genes related to P-bodies (A) Gene Ontology (GO) analysis of the downregulated genes upon knockdown of YAP/TAZ in HCT116 cells. The graph shows enrichment in the cellular component category. (B) ChIP–qPCR analysis of endogenous TEAD4 binding to the genomic locus of the indicated P-body-related genes in HCT116 cells. The CTGF promoter was included as the positive control. (C) qPCR analysis of the mRNA levels of the indicated P-body-related genes in YAP/TAZ knockdown HCT116, MDA-MB-231 and A549 cells. Cells were transfected with YAP/TAZ siRNA for 3 days before qPCR analysis. (D) qPCR analysis of the mRNA levels of the indicated P-body-related genes in HCT116, MCF7 and A549 cells stably expressing YAP^5SA^ and YAP^5SA-S94A^. Cells were infected with YAP^5SA^- and YAP^5SA-S94A^-containing retroviruses and selected with puromycin for one week before qPCR analysis. n=3 biologically independent samples per group. Two-tailed Student’s t test (B) and one-way ANOVA (C, D) were performed to assess statistical significance in this figure. These data (B-D) are representative of 3 independent experiments.

### YAP/TAZ enhances P-body formation

In contrast to stress granule (SG) formation, P-body formation is constitutive and independent of the activation of the integrated stress response (ISR) (Luo et al, 2018; Riggs et al, 2020). DEAD-box ATP-dependent RNA helicase 6 (DDX6) and mRNA-decapping enzyme 1A (DCP1A) are the essential components of P-bodies and are normally used as the biomarkers for P-bodies (James et al, 2010; Lavalee et al, 2021; Luo et al, 2018). To explore whether YAP/TAZ regulate P-body formation, we performed immunofluorescence analysis of DDX6 and DCP1A in YAP/TAZ knockdown cells plated at a low density. We found that knockdown of YAP/TAZ significantly decreased but overexpression of YAP^5SA^ increased the number of DDX6/DCP1A-positive foci in HCT116 cells (Figure 2A, 2B). HCT116 cells expressing YAP^5SA-S94A^ and control HCT116 cells showed similar numbers of P-bodies, which indicated that the TEAD transcription factors mediate the enhanced effects of YAP/TAZ on P-body formation (Figure 2B). Similar results were observed in YAP/TAZ knockdown MDA-MB-231 cells and YAP^5SA^/YAP^5SA-S94A^- expressing A549 cells (Figure 2-Figure Supplement 1A, B).

**Figure 2.**
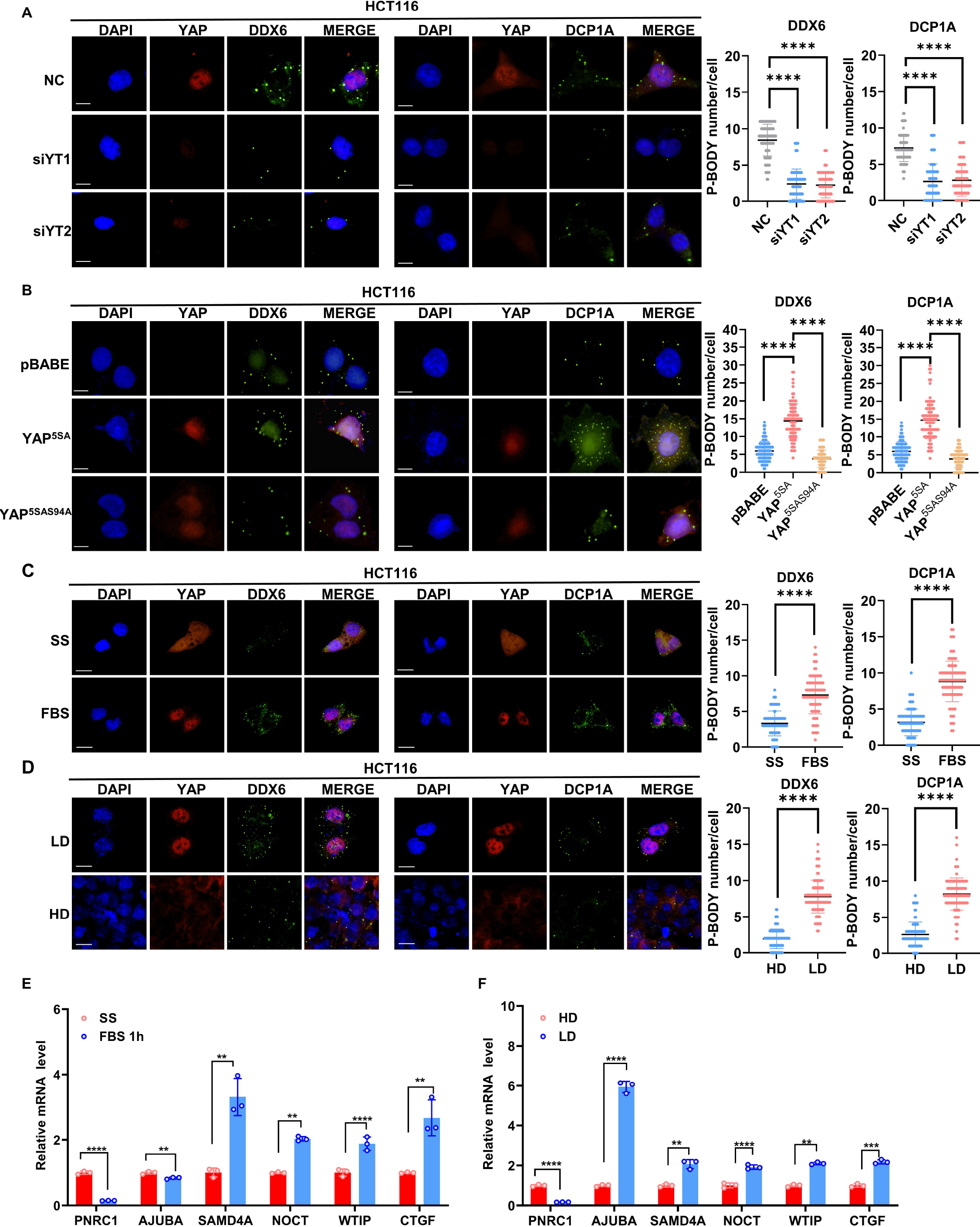
YAP/TAZ promotes P-body formation in CRC cells (A) Immunofluorescence analysis of the P-body markers DDX6 and DCP1A in YAP/TAZ knockdown HCT116 cells. Cells were transfected with YAP/TAZ siRNA for 3 days before processing for immunofluorescence staining using anti-DDX6 and anti-DCP1A antibodies. Foci were counted in 100 cells per group. (B) Immunofluorescence analysis of DDX6 and DCP1A in HCT116 cells expressing YAP^5SA^ _and YAP_5SA-S94A_._ (C) Immunofluorescence analysis of DDX6 and DCP1A in HCT116 cells. Cells were treated with 10% FBS for 1 h after overnight serum starvation (SS). (D) Immunofluorescence analysis of DDX6 and DCP1A in HCT116 cells in sparse or confluent culture. (E, F) qPCR analysis of the indicated genes in HCT116 cells. HCT116 cells were treated with 10% FBS for 1 h after overnight serum starvation (E) or culture under sparse or confluent conditions in standard culture medium (F). Kruskal-Wallis test (A, B), Mann– Whitney U test (C, D) and Two-tailed Student’s t test (E, F) were performed to assess statistical significance. These data (A-F) are representative of 3 independent experiments.

YAP/TAZ are well known to be activated by serum stimulation and suppressed by high cell densities (Yu et al, 2012; Zhao et al, 2007). Of note, cytoplasmic translocation of YAP at high cell density was first observed in the untransformed NIH3T3 cells (Zhao et al, 2007). Thus, in addition to a series of cancer cell lines, NIH3T3 cells were further included in this study. Consistently, overexpression of YAP^5SA^ but not the YAP^5SA-S94A^ increased the number of DDX6/LSM14A-positive foci in NIH3T3 cells (Figure 2-Figure Supplement 2A).

Upregulation of *Ajuba, Samd4* (mouse ortholog of human SAMD4A) and *Noct* and downregulation of *Pnrc1* was also observed in NIH3T3 cells overexpressed with YAP^5SA^ but not cells with YAP^5SA-S94A^ overexpression (Figure 2-Figure Supplement 2E). Moreover, despite their constitutive formation in cells, the size and number of P-bodies are altered in response to stress (Luo et al, 2018; Riggs et al, 2020). Next, we evaluated P-bodies under exposure to different stimuli. We observed that serum stimulation led to rapid induction of P-body formation in HCT116 and NIH3T3 cells (Figure 2C, Figure 2-Figure Supplement 2B). Knockdown of YAP/TAZ attenuated the enhancement of P-body formation induced by serum stimulation (Figure 2-Figure Supplement 1C). Conversely, at a high cell density, the number of P-bodies was significantly decreased in HCT116 and NIH3T3 cells (Figure 2D, Figure 2-Figure Supplement 2C). Consistent with this finding, the expression of SAMD4A, NOCT, and WTIP in HCT116 cells was induced by serum stimulation and suppressed by culture at a high cell density (Figure 2E, 2F). Similar results were also observed in NIH3T3 cells (Figure 2-Figure Supplement 2F, Figure 2-Figure Supplement 2G). Intriguingly, both serum starvation and culture at a high cell density dramatically increased the expression of PNRC1, consistent with the tumor suppressor function of PNRC1 (Figure 2E, 2F, Figure 2-Figure Supplement 2F, Figure 2-Figure Supplement 2G). Recent studies have revealed that mechanical cues as an important signal modulating YAP/TAZ activity (Aragona et al, 2013; Dupont et al, 2011). Diverse mechanical forces, such as increased ECM rigidity, cell stretching, shear stress, or the increased area of cell adhesion, can all activate YAP, which is dominant over Hippo signaling (Dasgupta & McCollum, 2019; Piccolo et al, 2014). Next, we examined whether ECM stiffness affected P-body formation. When NIH3T3 cells were shifted from soft (1kPa) to stiff (40kPa) matrices, YAP was translocated into nucleus and activated (Figure 2-Figure Supplement 2D). Furthermore, the P-body formation was enhanced which was associate with decreased mRNA level of *Pnrc1* and increased mRNA levels of *Ajuba, Samd4* and *Noct* (Figure 2-Figure Supplement 2D, Figure 2-Figure Supplement 2H). Collectively, our data indicate that YAP/TAZ could be positive regulators of P-body formation in response to various stimuli, probably by modulating the expression of P-body-related genes.

### SAMD4A, AJUBA and PNRC1 mediate the functions of YAP/TAZ in regulating P-body formation

Next, we investigated whether the P-body-related genes transcriptionally regulated by YAP/TAZ mediate the biological functions of YAP/TAZ in regulating P-body formation. The LIM-domain proteins AJUBA, WTIP and LIMD1 are known as negative regulators of LATS1 (Das Thakur et al, 2010). They are also components of P-bodies and are required for miRNA-mediated silencing (James et al, 2010). SAMD4A is the mammalian homolog of Drosophila Smaug, which is involved in translational repression and localized in P-bodies (Baez & Boccaccio, 2005). First, we knocked down AJUBA and SAMD4A in HCT116 cells overexpressing YAP^5SA^ (Figure 3-Figure Supplement 1A, B). As expected, knockdown of both AJUBA and SAMD4A significantly diminished the promoting effect of YAP^5SA^ overexpression on P-body formation in HCT116 cells (Figure 3A). Unlike AJUBA and SAMD4A, PNRC1 is a tumor suppressor that inhibits P-body formation by recruiting cytoplasmic DCP1A/DCP2 into the nucleolus, thus loss of cytoplasmic DCP1A/DCP2 results in disruption of P-body (Gaviraghi et al, 2018). Overexpression of YAP^5SA^ suppressed PNRC1 expression; thus, WT PNRC1 and PNRC1 with the W300A mutation, which disrupts the interaction between PNRC1 and DCP1A/DCP2, were overexpressed in YAP^5SA^-expressing HCT116 cells (Figure 3-Figure Supplement 1C, D) (Gaviraghi et al, 2018). We observed that overexpression of WT PNRC1 but not the W300A mutant dramatically decreased the number of P-bodies in YAP^5SA^-expressing HCT116 cells (Figure 3B). We also examined whether the attenuation of P-body formation by knockdown of YAP/TAZ can be restored by knockdown of PNRC1 (Figure 3-Figure Supplement 1E). Consistent with the above findings, the reduction in the P-body number was reversed by knockdown of PNRC1 in YAP/TAZ knockdown HCT116 cells (Figure 3-Figure Supplement 1F). Collectively, these findings indicate that YAP/TAZ enhance P-body formation through modulation of a series of P-body-related genes. Both activation of SAMD4A and AJUBA expression and downregulation of PNRC1 are involved in YAP/TAZ-induced P-body formation.

**Figure 3.**
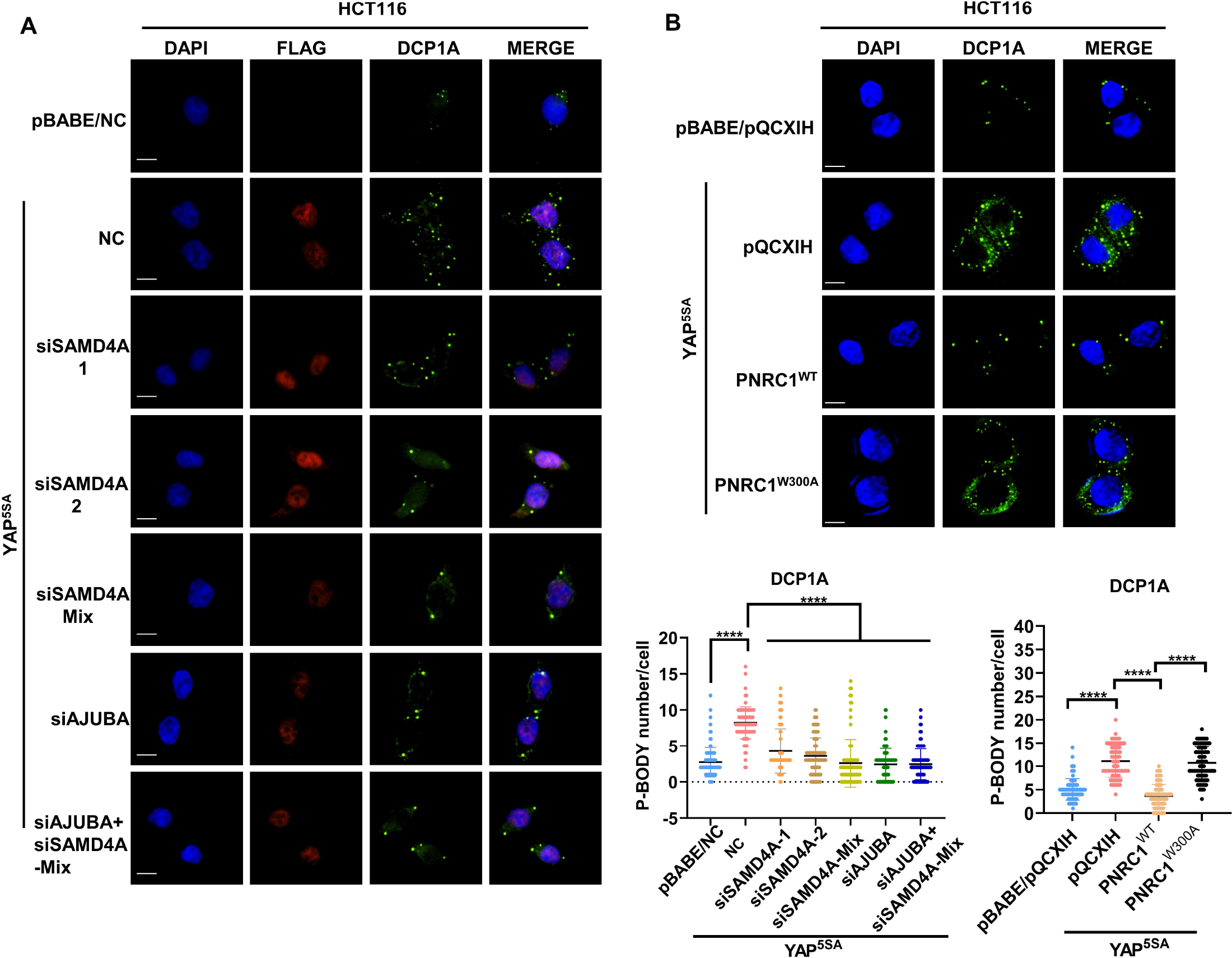
SAMD4A, AJUBA, and PNRC1 mediate the regulatory functions of YAP/TAZ in P-body formation (A) Immunofluorescence analysis of DDX6 and DCP1A in HCT116 cells stably expressing YAP^5SA^ and YAP^5SA^-expressing cells transiently transfected with SMAD4A and AJUBA siRNA. Foci were counted in 100 cells per group. (B) Immunofluorescence analysis of DDX6 and DCP1A in HCT116 cells expressing YAP^5SA^ alone or in combination with PNRC1^WT^ or PNRC1^W300A^. Kruskal-Wallis test was performed to assess statistical significance. These data (A-B) are representative of 3 independent experiments.

### YAP/TAZ inhibit PNRC1 gene transcription by recruiting the NuRD complex

PNRC1 is a newly identified tumor suppressor gene whose expression is frequently downregulated in cancer (Gaviraghi et al, 2018). Thus, we further explored the molecular mechanism of YAP/TAZ in inhibiting the PNRC1 gene transcription. The ChIP-seq data for TEAD4 at the PNRC1 gene locus in multiple cancer cells implicated PNRC1 as a potential direct target gene of YAP/TAZ-TEAD transcription complexes (Figure 1-Figure Supplement 1B). As ChIP–qPCR analysis of TEAD4 in HCT116 cells revealed one TEAD4 binding site at the PNRC1 promoter and another in the PNRC1 intron, we constructed PNRC1 promoter and PNRC1 intron luciferase reporter plasmids. We observed that overexpression of YAP^5SA^ significantly decreased the luciferase activity of both the PNRC1 promoter and intron reporters (Figure 4A). Compared with the 5SA mutation in YAP, the S94A mutation resulted in a decreased suppressive effect on PNRC1 promoter and intron luciferase reporter activity (Figure 4A). In contrast, the luciferase activity of both the PNRC1 promoter and intron reporters was significantly enhanced in YAP/TAZ knockdown HCT116 cells (Figure 4B, Figure 4-Figure Supplement 1A). Bioinformatic analysis of TEAD4 ChIP peaks in the PNRC1 promoter and intronic regions with JASPAR revealed the existence of one TEAD binding motif in each peak region; thus, we further constructed PNRC1 luciferase reporter plasmids with mutated TEAD binding sites. Consistent with the above results, mutation of the TEAD binding sites abolished the inhibitory effect of YAP^5SA^on the PNRC1 promoter and intron luciferase reporters (Figure 4C). Similarly, mutation of the TEAD binding sites escaped the derepression of PNRC1 promoter and intron luciferase reporters by YAP/TAZ knockdown (Figure 4D). Furthermore, the ChIP–qPCR results confirmed that YAP bound to the promoter and intronic regions of PNRC1, which required its interaction with TEADs (Figure 4-Figure Supplement 1B, Figure 4-Figure Supplement 1C).

**Figure 4.**
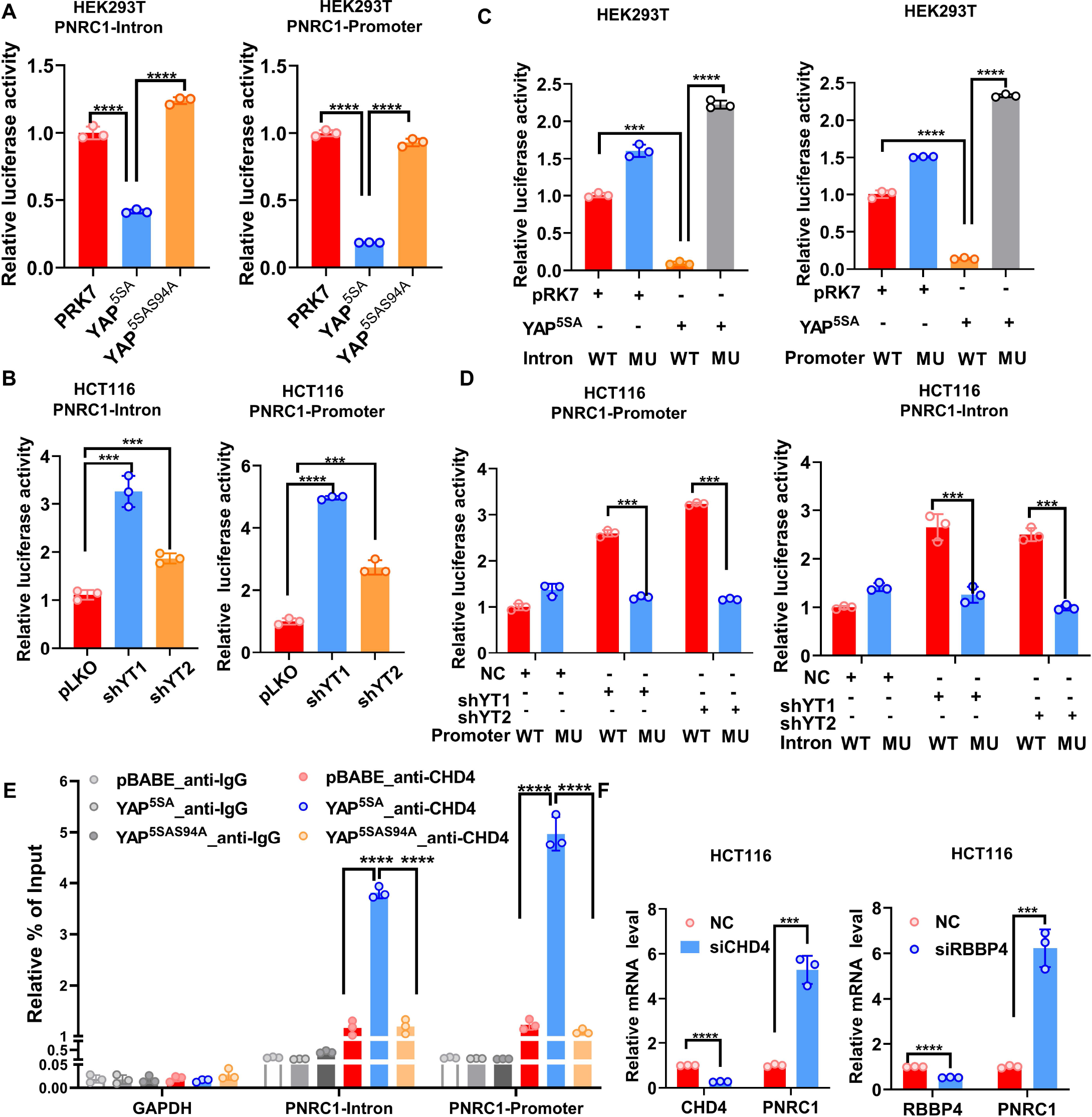
YAP suppresses PNRC1 gene transcription by recruiting the NuRD complex (A) Overexpression of YAP^5SA^ but not YAP^5SA-S94A^ decreased the luciferase activity of the PNRC1 promoter and intron reporters. HEK-293T cells were transfected with the indicated FLAG-YAP^5SA^ and YAP^5SA-S94A^ expression plasmids and the PNRC1 promoter or intron luciferase reporter. (B) Knockdown of YAP/TAZ stimulated the luciferase activity of the PNRC1 promoter and intron reporters. The PNRC1 promoter or intron luciferase reporter plasmid and the Renilla luciferase reporter plasmid were cotransfected into HCT116 cells stably expressing pLKO- vec, shYAP/TAZ-1, or shYAP/TAZ-2. (C, D) Luciferase assay of the PNRC1 promoter/intron WT reporters and mutant reporters with TEAD binding motif mutations in HEK-293T cells (C) and HCT116 cells (D). (E) ChIP–qPCR analysis of CHD4 binding to the PNRC1 promoter and intronic regions in control and HCT116 cells stably expressing FLAG-YAP^5SA^ or YAP^5SA-S94A^. (F) qPCR analysis of PNRC1, CHD4, and RBBP4 in HCT116 cells transfected with the indicated siRNAs. n=3 biologically independent samples per group. One-way ANOVA (A- E) and two-tailed Student’s t test (F) were performed to assess statistical significance in this figure. These data (A-F) are representative of 2 independent experiments.

In addition to functioning as transcriptional coactivators, YAP/TAZ can also act as transcriptional corepressors by recruiting the NuRD complex (Kim et al, 2015b). The ChIP–qPCR results showed that the NuRD complex component CHD4 was recruited to the promoter and intronic regions of the PNRC1 gene by overexpressed YAP^5SA^ but not by the TEAD binding-defective YAP^5SA-S94A^ mutant (Figure 4E). Compared to the genomic locus of PNRC1, the binding enrichment of CHD4 at the YAP target genes activated by YAP/TAZ were relative lower and wasn’t affected by overexpression of YAP (Figure 4-Figure Supplement 1D). Moreover, knockdown of the NuRD complex components CHD4 and RBBP4 significantly upregulated the mRNA expression of PNRC1 in HCT116 cells (Figure 4F). Consistently, knockdown of CHD4 significantly decreased the number of DDX6/DCP1A-positive foci in HCT116 cells (Figure 4-Figure Supplement 1E). Taken together, these data demonstrate that YAP/TAZ inhibit PNRC1 gene transcription through direct binding of TEADs to the PNRC1 gene locus and that the NuRD complex is required for the transcriptional repression of PNRC1 by YAP/TAZ.

### PNRC1 suppresses the oncogenic function of YAP in CRC

Analysis of colorectal (COAD) and rectal (READ) TCGA datasets revealed that the mRNA level of PNRC1 was significantly decreased in CRC (Figure 5-Figure Supplement 1A). We further confirmed the decreased mRNA level of PNRC1 in CRC by qPCR analysis of 16 CRC tissues with paired normal mucosal tissues; this finding implies that PNRC1 is a potential tumor suppressor also in CRC (Figure 5-Figure Supplement 1B). Thus, we sought to explore whether downregulation of PNRC1 mediates the oncogenic function of YAP in CRC. To this end, we examined whether the YAP overexpression-induced oncogenic phenotype can be attenuated by coexpression of YAP^5SA^ with WT PNRC1 or the W300A mutant in HCT116 cells. We observed that reexpression of WT PNRC1 almost completely abolished the increases in cell proliferation and colony formation induced by YAP^5SA^ overexpression in HCT116 cells (Figure 5A, 5B). Re-expression of the PNRC1 W300A mutant did not affect the proliferation and colony formation of YAP^5SA^-expressing HCT116 cells, which implied that the suppressive effect of PNRC1 on YAP relies on the recruitment of cytoplasmic DCP1A/DCP2 into the nucleolus by PNRC1 (Figure 5A, 5B). Similarly, overexpression of PNRC1 WT but not PNRC1 W300A suppressed the increase in migration induced by YAP^5SA^ in HCT116 cells (Figure 5C). To verify the tumor-suppressive effect of PNRC1 on YAP in CRC *in vivo*, we performed a xenograft assay by subcutaneously injecting HCT116 cells into nude mice. Consistent with the above findings, reexpression of WT PNRC1 but not the W300A mutant dramatically inhibited the growth of YAP^5SA^-expressing HCT116 xenografts, and xenograft tumors formed from HCT116 cells coexpressing YAP^5SA^ and PNRC1 were significantly smaller than tumors formed from HCT116 cells expressing YAP^5SA^ alone or in combination with the PNRC1 W300A mutant (Figure 5D). Ki67 staining of xenograft tumors further showed fewer Ki67-positive cells in xenograft tumors formed from HCT116 cells coexpressing YAP^5SA^ and PNRC1 (Figure 5E, Figure 5-Figure Supplement 1C). Next, we examined whether the YAP/TAZ knockdown-induced attenuation of the oncogenic phenotype can be restored by knockdown of PNRC1 in HCT116 cells. Intriguingly, the decrease in proliferation and attenuation of migration induced by YAP/TAZ knockdown were reversed by knockdown of PNRC1 in HCT116 cells (Figure 5-Figure Supplement 1D, E). Overall, these results indicate that YAP promotes tumorigenesis by downregulating PNRC1 expression and that reexpression of PNRC1 suppresses YAP-driven tumor growth.

**Figure 5.**
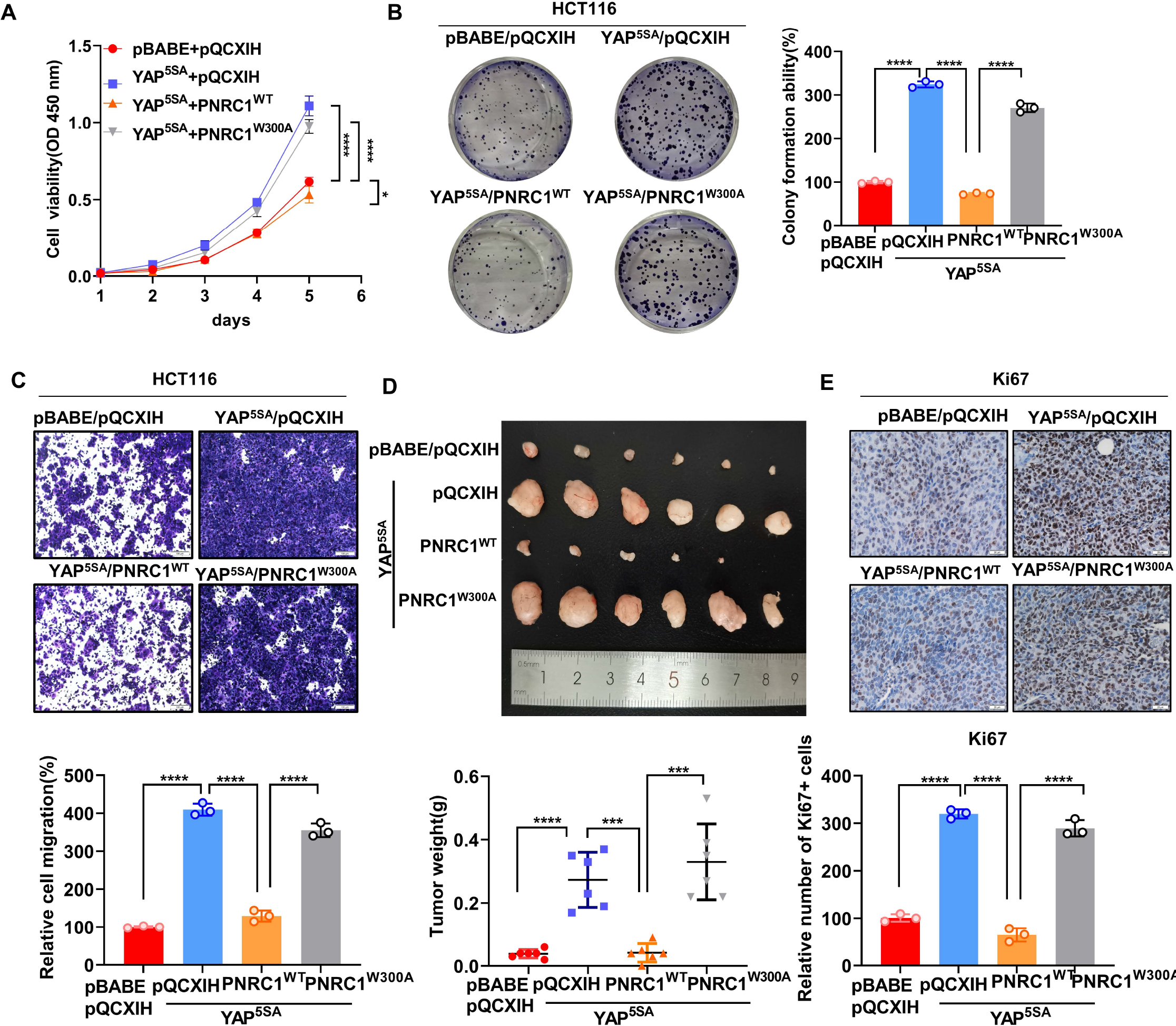
PNRC1 attenuates the oncogenic function of YAP in CRC (A) CCK8 proliferation assays of HCT116 cells stably expressing YAP^5SA^ alone or in combination with of PNRC1^WT^ or PNRC1^W300A^. n=5 biologically independent samples per group. (B, C) Colony formation assay (B) and Transwell assay (C) of HCT116 cells stably expressing YAP^5SA^ alone or in combination with PNRC1^WT^ or PNRC1^W300A^. n=3 biologically independent samples per group. (D) Representative images of xenograft tumors formed from HCT116 cells stably expressing YAP5SA alone or in combination with of PNRC1^WT^ or PNRC1^W300A^ (n = 6). (E) Representative images of IHC staining of the proliferation marker Ki67 in xenograft tumors formed from HCT-116 cells stably expressing YAP^5SA^ alone or in combination with PNRC1^WT^ or PNRC1^W300A^ (n = 3). Two-way ANOVA (A) and one-way ANOVA (B-E) were performed to assess statistical significance in this figure. These data (A-C) are representative of 2 independent experiments.

### P-body disassembly attenuates YAP-driven cell proliferation and migration in CRC

Due to the inhibitory effects of PNRC1 on P-body formation and the YAP-induced oncogenic phenotype, we evaluated whether enhanced P-body formation plays a vital role in YAP-driven cancer cell proliferation and migration. The proteins LSM14 homolog A (LSM14A) and DDX6 are essential nucleating proteins for P-body assembly, and DCP1A plays a vital role in further RNP aggregation, which is required for stress-dependent P-body aggregation (Lavalee et al, 2021; Luo et al, 2018; Riggs et al, 2020). To explore the requirement of P-body formation for the YAP-induced oncogenic phenotype, we generated YAP^5SA^-expressing HCT116 cell lines with stable knockdown of DCP1A, LSM14A and DDX6 (Figure 6-Figure Supplement 1A, Figure 6-Figure Supplement 2A-C). By immunofluorescence analysis of DDX6 and DCP1A, we further confirmed the knockdown of DCP1A and DDX6 and observed a reduced number of P-bodies upon knockdown of DCP1A, LSM14A or DDX6 in YAP^5SA^-expressing HCT116 cells (Figure 6-Figure Supplement 1B). Next, by using a CCK8 assay, we found that knockdown of DCP1A, LSM14A and DDX6 suppressed the proliferation of YAP^5SA^-expressing and control “parental” HCT116 cells, consistent with the results of the colony formation assay (Figure 6A, 6B). As an oncogene, YAP is known to promote cell division and inhibit cell apoptosis of cancer cells (He et al, 2020; Jang et al, 2017). By analyzing the cell cycle and cell apoptosis, we further found that knockdown of DCP1A, LSM14A and DDX6 all led to downregulation of cell mitosis and increased cell apoptosis, which was opposite to the effect of YAP^5SA^ overexpression in HCT116 cells (Figure 6-Figure Supplement 3A-B). Furthermore, knockdown of either DCP1A or LSM14A significantly attenuated the enhancement of cell migration induced by overexpression of YAP^5SA^ in HCT116 cells (Figure 6C). In contrast, knockdown of DDX6 showed stimulative effect on the migration of both control and YAP^5SA^-expressing HCT116 cells, possibly due to the diverse functions of DDX6 (Figure 6-Figure Supplement 4A) (Di Stefano et al, 2019). To further demonstrate the potential role of P-body mediating the function of YAP/TAZ in CRC, we established YAP^5SA^-expressing HCT116 cell lines with stable knockdown of AJUBA and SAMD4A (Figure 6-Figure Supplement 4B, C). Indeed, both knockdown of AJUBA and SAMD4A suppressed the proliferation and cell migration of YAP^5SA^-expressing and control “parental” HCT116 cells (Figure 6-Figure Supplement 4D, E). Collectively, our data demonstrate that P-body formation is required for the oncogenic function of YAP in CRC.

**Figure 6.**
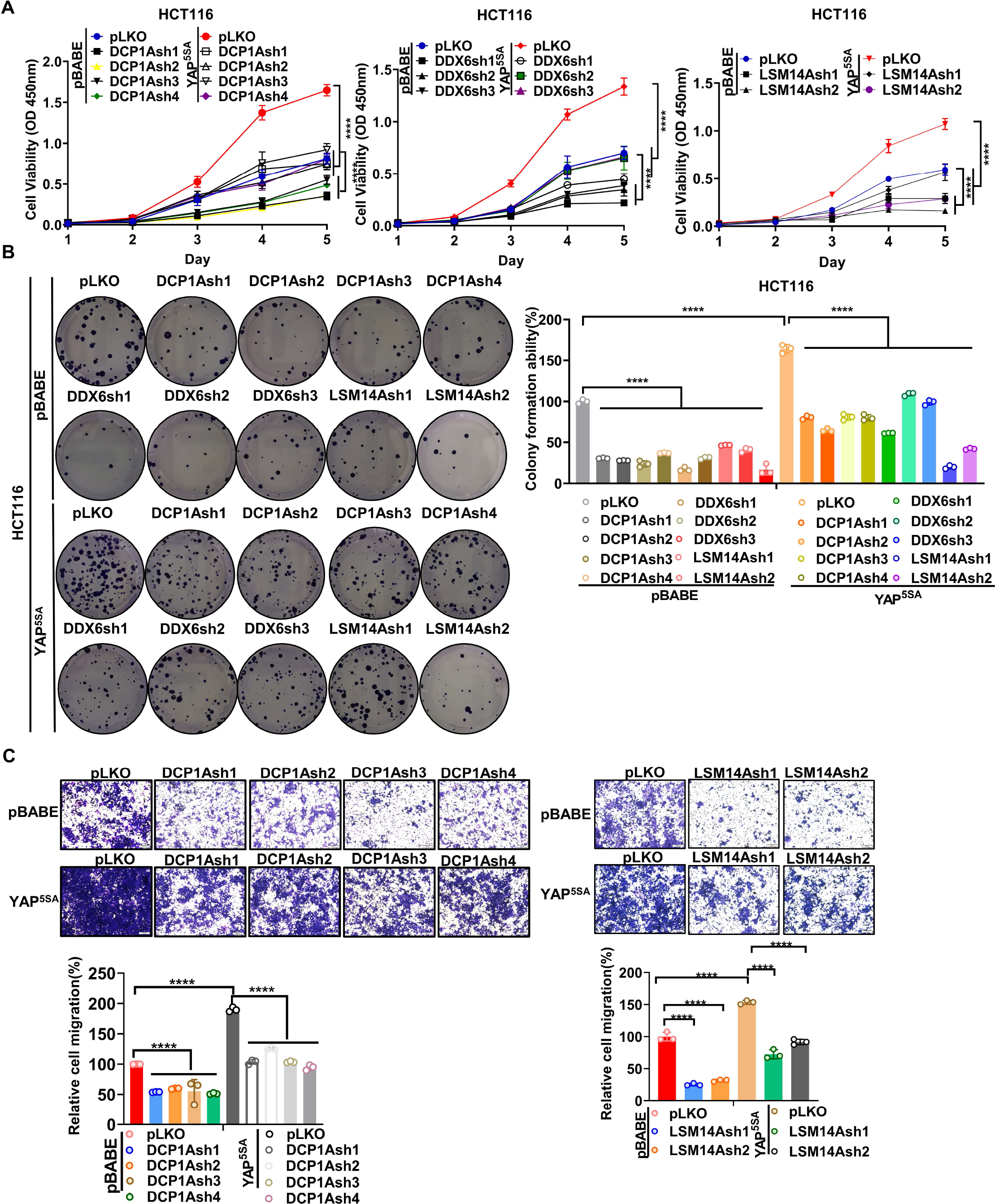
Knockdown of P-body-related core genes suppresses the oncogenic function of YAP in CRC (A) CCK8 proliferation assays of control HCT116 cells with or without knockdown of DCP1A, LSM14A or DDX6 and HCT116 cells stably expressing YAP^5SA^ with or without knockdown of DCP1A, LSM14A or DDX6. n=5 biologically independent samples per group. (B, C) Colony formation assay (B) and Transwell assay (C) of control HCT116 cells with or without knockdown of DCP1A, LSM14A or DDX6 and HCT116 cells stably expressing YAP^5SA^ with or without knockdown of DCP1A, LSM14A or DDX6. n=3 biologically independent samples per group. Two-way ANOVA (A) and one-way ANOVA (B, C) were performed to assess statistical significance in this figure. These data (A-C) are representative of 3 independent experiments.

### Codependency of YAP/TEAD and essential P-body-related genes in pancancer CRISPR screens

Based on the observation that P-body disassembly attenuates YAP-driven cell proliferation in CRC cells, we speculated that cancer cells whose proliferation is dependent on YAP should also be vulnerable to knockout of essential P-body genes. To this end, we analyzed the Cancer Dependency Map (DepMap), which aims to systematically assess the effect of single-gene inactivation on cell proliferation by CRISPR and shRNA screens and define genetic dependencies in hundreds of cancer cell lines by integrating data pertaining to multiple molecular characteristics, such as Cancer Cell Line Encyclopedia (CCLE) data (Dempster et al, 2021; Tsherniak et al, 2017). As expected, by analyzing gene expression data from the CCLE, we observed a strong positive correlation between YAP-regulated P-body-related genes (SAMD4A, AJUBA, and WTIP) and canonical target genes of YAP (CTGF, CYR61, AXL, and AMOTL2) in cell lines across cancers or in cell lines of colorectal, breast and lung lineages (Figs. 7A, Figure 7-Figure Supplement 1A, Figure 7-Figure Supplement 1B). IHC analysis of 294 CRC tissues further showed the positive correlation between the expression of AJUBA/SAMD4A and YAP (Figure 7-Figure Supplement 1D). Although there were no correlations between PNRC1 and YAP target genes in cell lines across cancers, we found that the mRNA level of PNRC1 was negatively correlated with that of YAP target genes in cancer cells of thyroid and central nervous system (CNS) lineages (Figure 7B, Figure 7-Figure Supplement 1C). Next, we analyzed the effect of P-body core gene knockout in 1070 cancer cell lines (DepMap 22Q1 Public+Score, Chronos). Strikingly, knockout of the P-body -nucleation-determining genes DDX6 and LSM14A inhibited proliferation in multiple cancer cell lines (negative Chronos score) (Figure 7C). Similar results were observed for EDC4, which is required for P-body aggregation (Figure 7C). Logically, correlations between dependency profiles suggests functionality in the same pathway or regulatory axis; thus, EDC4 is strongly associated with multiple known P-body genes (DDX6, DCP2, EIF4ENIF1, etc.) (Appendix Table S2). Furthermore, we found that YAP ranked 14th among genes correlating with EDC4 (Figure 7C, Appendix Table S2). In addition, the YAP dependency score was positively correlated with the DDX6 and LSM14A scores (Figure 7C). Similar relationships were observed between EDC4/DDX6/LSM14A and TEAD1/3 (Figure 7-Figure Supplement 2). Last, we examined whether cell proliferation and cell migration are affected by the knockdown of DCP1A or LSM14A and overexpression of PNRC1 in MCF7, MDA-MB-231 and A549 cell lines, whose proliferation are dependent on YAP/TAZ activity. In consistent to the observation in HCT116 cells, the knockdown of DCP1A/LSM14A and overexpression of PNRC1 attenuated both cell proliferation and cell migration in these three YAP-dependent cancer cells (Figure 7- Figure Supplement 3-5). Collectively, the codependencies of YAP/TEAD and essential P-body genes further suggests that enhanced P-body formation plays a vital role in YAP- induced tumorigenesis.

**Figure 7.**
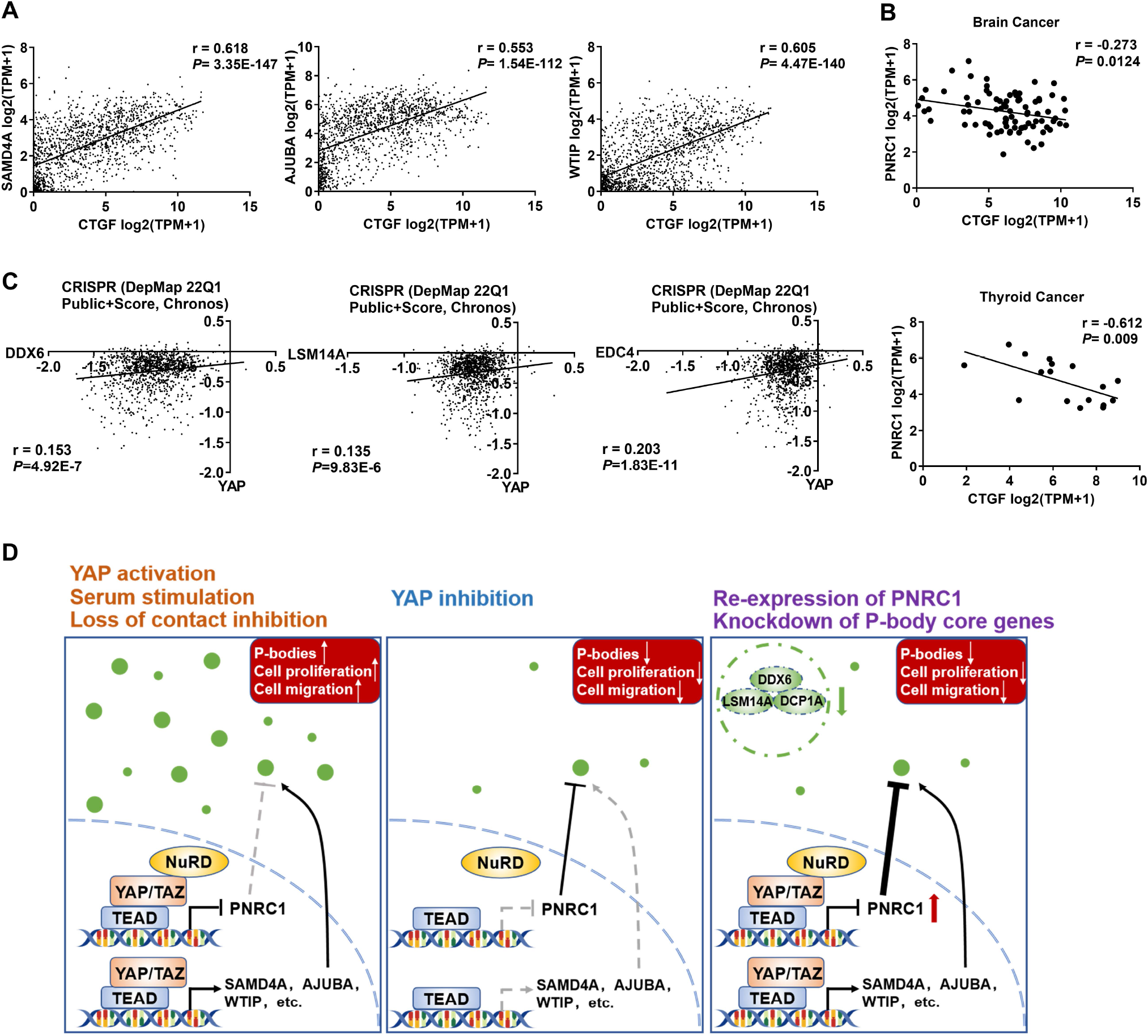
DepMap analysis reveals the codependencies of YAP/TEAD and P-body core genes in pancancer CRISPR screens (A) Positive correlations between the mRNA levels of CTGF and SAMD4A/AJUBA/WTIP in 1393 cancer cell lines. (B) Negative correlation between the mRNA levels of CTGF and PNRC1 in brain cancer cell lines (n=83) and thyroid cancer cell lines (n=17). (C) Positive correlations between the dependency scores of YAP and DDX6/LSM14A/EDC4 in 1070 cancer cell lines. The Chronos dependency scores were extracted from the DepMap database. The negative Chronos score indicates decreased cell proliferation upon gene knockout. Pearson correlation analysis was used to assess statistical significance. (D) In response to serum stimulation or under loss of contact inhibition or reduced ECM stiffness, activation of YAP enhances the P-body formation to promote CRC cell proliferation and migration. Disruption of P-bodies by overexpression of the tumor suppressor gene PNRC1 or knockdown of P-body core genes could attenuate the cell proliferation and migration induced by activation of YAP in CRC cells.

## Discussion

Dysregulation of the Hippo pathway occurs in a variety of cancers, leading to cell transformation and diverse changes in tumor cells through activation of the YAP/TEAD transcriptional program (Calses et al, 2019; Kulkarni et al, 2020; Nguyen & Yi, 2019; Wang et al, 2018; Zanconato et al, 2016). Here, we demonstrated a crucial role of YAP/TEAD in regulating P-body formation in multiple cancer cell lines. Through transcriptional stimulation of positive regulators of P-body formation (AJUBA, WTIP, and SAMD4A) and suppression of negative regulators of P-body formation (PNRC1), YAP enhances P-body formation and increases the number of P-bodies in cancer cells, which suggests that YAP is a positive regulator of P-body formation (Figure 7D). Studies of P-bodies in yeast have shown that the size and number of P-bodies increase upon exogenous and endogenous stress (Luo et al, 2018). In addition, our study revealed that the number of P-bodies decreases under serum starvation, contact inhibition and decreased ECM rigidity, possibly due to inactivation of YAP/TEAD. In contrast, a recent study, which provided the first link between YAP and P-bodies, implicated YAP as a negative regulator of P-bodies in KHSV-infected HUVECs (Castle et al, 2021). Elizabeth L. Castle et al. reported that virus-encoded Kaposin B (KapB) induces actin stress fiber formation and disassembly of P-bodies, which requires RhoA activity and the YAP transcriptional program (Castle et al, 2021). YAP- enhanced autophagic flux was proposed to participate in KapB-induced P-body disassembly, consistent with the concept that stress granules and P-bodies are cleared by autophagy (Buchan et al, 2013; Castle et al, 2021). However, an increasing number of studies have reported the contradictory role of YAP in autophagy regulation, which suggests that YAP-mediated autophagy regulation is cell type- and context-dependent (Jin et al, 2021; Pei et al, 2022; Totaro et al, 2019; Wang et al, 2020). Furthermore, though YAP is required for the cell proliferation in HUVEC, transformed cell lines often display elevated baseline YAP/TAZ activity compared to normal cells and possess many alterations in growth signaling pathways including autophagy signaling (Nguyen & Yi, 2019; Shen & Stanger, 2015; Zanconato et al, 2016). Thus, the contradictory observations regarding the role of YAP in modulating P-body formation between Elizabeth L. Castle et al.’s study and our study could be due to the different cell contexts and different cell conditions (baseline vs. KHSV infection).

In addition to the transcriptional regulation, P-body dynamics can also be modulated by post-translational modifications (Luo et al, 2018). P-body constituent proteins, such as DCP1A and DCP2, are phosphorylated, which affects the protein interaction between DCP1A and DCP2 and subsequent P-body assembly (Chiang et al, 2013; Yoon et al, 2010). Previous studies have shown that DCP1A is hyperphosphorylated during mitosis and P-body assembly is dynamically changed across the cell cycle (Aizer et al, 2013; Yang et al, 2004). Meanwhile, Hippo signaling is intrinsically regulated and YAP can also be directly phosphorylated by CDK1 during cell cycle progression (Kim et al, 2019; Yang et al, 2013). Besides, as a direct regulator of P-body formation, AJUBA is also phosphorylated by CDK1 and mitotic phosphorylation of AJUBA promotes cancer cell proliferation (Chen et al, 2016). Thus, we speculated that mitotic phosphorylation of YAP and AJUBA might also play a potential role in modulating P-body dynamics during cell cycle.

Compared with the role of stress granules, the role of P-bodies in tumorigenesis and tumor progression is not well studied and is considered to be cancer type- or context-dependent (Lavalee et al, 2021). TGF-β induces P-body formation and epithelial-mesenchymal transition (EMT) in mammary epithelial cells, while inhibition of P-body formation by knockdown of DDX6 reverses EMT and suppresses breast cancer metastasis, implying a prometastatic function of P-bodies during the progression of breast cancer (Hardy et al, 2017). In prostate cancer cells, dephosphorylation of EDC3 promotes the localization of EDC3-containing P-bodies and increases the P-body number (Bearss et al, 2021). The increase in EDC3-containing P-bodies leads to sequestration or decay of a subset of mRNAs related to cell attachment and cell growth, such as ITGB1, ITGA6 and KLF4, which ultimately inhibits cell proliferation and cell migration (Bearss et al, 2021). Of note, EDC3 and LSM14A compete for binding to the P-body core protein DDX6, and P-body formation still occurs constitutively in EDC3 KO prostate cancer cells (Bearss et al, 2021). Therefore, it can be speculated that there might be different types of P-bodies that contain different RNAs and exert protumorigenic or tumor-suppressive functions in different cell contexts. In CRC, DCP1A expression is elevated, which is associated with advanced TNM stages, lymph node metastasis and poor prognosis, and overexpression of DCP1A enhances P-body formation (Wu et al, 2018a; Wu et al, 2018b). These studies imply the potential protumorigenic function of P-bodies in CRC. Furthermore, our study showed that disruption or attenuation of P-body formation by knockdown of YAP-regulated P-body-related genes or the P-body core genes (DDX6, DCP1A, LSM14A) suppressed YAP-induced oncogenic phenotypes in CRC cells, such as cell proliferation and cell migration, further indicating the protumorigenic function of P-bodies in CRC or at least in CRC with active YAP. Numerous studies have demonstrated the YAP/TAZ promote cancer cell growth through directly transcriptional regulation of genes related with cell cycle and cell apoptosis (He et al, 2020; Jang et al, 2017). Since P-bodies control the storage of untranslated mRNAs, YAP/TAZ might modulate gene expression through indirectly promoting P-body formation and the storage of untranslated target mRNAs. Future work is needed to explore P-body-enriched RNAs in CRC cells, which will further uncover the underlying mechanism by which P-bodies mediate the oncogenic function of YAP.

Recently, a study exploring new TSGs based on hemizygous deletions in multiple cancers revealed that PNRC1 is a novel tumor suppressor gene (Gaviraghi et al, 2018). PNRC1 translocates the cytoplasmic DCP1A/DCP2 decapping complex into the nucleolus, which subsequently impedes rRNA transcription and ribosome biogenesis (Gaviraghi et al, 2018). This translocation of DCP1A/DCP2 also leads to disassembly of P-bodies; thus, PNRC1 could also inhibit tumor cell proliferation by disrupting P-body formation. Moreover, hemizygous deletion of the 6q15 locus, where PNRC1 is located, occurs in multiple cancers, including prostate, pancreatic, breast and liver cancers (Gaviraghi et al, 2018).

Our findings suggest that transcriptional downregulation of PNRC1 by YAP activation could be a new mechanism of PNRC1 dysregulation during tumorigenesis. In addition, multiple oncogenes, such as MYC, RAS and PI3K, can activate rRNA transcription and boost ribosome biogenesis to support cancer cell proliferation (Pelletier et al, 2018). The presence of the YAP-PNRC1 regulatory axis implies a potential role of YAP in ribosome biogenesis, which warrants further investigation in follow-up studies.

It has been shown that PNRC1 inhibits RAS- and MYC-driven tumor cell proliferation (Gaviraghi et al, 2018). In addition, YAP acts downstream of mutant KRAS, and activation of YAP was found to drive KRAS-independent tumor relapse in preclinical models of pancreatic cancer (Kapoor et al, 2014; Shao et al, 2014; Zhang et al, 2014). Strikingly, reexpression of PNRC1 also dramatically diminished the cell proliferation induced by YAP overexpression in CRC cells in our study. These data indicate that PNRC1 is a tumor suppressor gene with a strong antitumor effect on various oncogenes; thus, reexpression of PNRC1 could be a promising anticancer therapeutic strategy. In addition, as an alternative to targeting the YAP/TEAD complex, drugs that inhibit downstream effectors of YAP/TAZ have shown efficacy in the clinic (Gay et al, 2017; Neesse et al, 2013; Nguyen & Yi, 2019). The identification of the P-body as a new downstream effector of YAP/TAZ suggests that disruption of P-bodies might be a potential therapeutic strategy for tumors with active YAP. Although each P-body core gene performs multiple biological functions, unbiased functional CRISPR screening across cancer cell lines (DepMap) revealed that LOF of a series of P-body core genes significantly suppresses proliferation in various tumor cell lines. The functional overlap in P-body assembly and the positive correlation between the dependency profiles of these P-body core genes imply the important role of P-bodies in tumor cell proliferation and cell survival. Several compounds, including translation inhibitors, have been reported to inhibit P-body formation (Martinez et al, 2013; Stribinskis & Ramos, 2007). Notably, actin polymerization can activate YAP (Sun & Irvine, 2016). Methyl-chivosazol, an actin polymerization inhibitor, was found to be a strong inhibitor of P-body formation by screening of a library of compounds derived from myxobacteria (Martinez et al, 2013). However, these small molecules indirectly target P-bodies and show extensive effects on cells (Martinez et al, 2013; Stribinskis & Ramos, 2007). Thus, the development of inhibitors directly targeting P-body core proteins will provide a chemical tool for exploring the function of P-bodies in tumors and assess the therapeutic efficacy of P-body disassembly in cancer. Overall, our study reveals the P-body as a new downstream effector of YAP/TAZ, which opens a new possibility of targeting P-body assembly to combat tumors (Figure 7D).

## Materials and Methods

### Cell culture and transfection

HEK293T, NIH3T3, HCT116, MCF7, MDA-MB-231, and A549 cells were purchased from the American Type Culture Collection (ATCC) and authenticated by short tandem repeat analysis. HEK293T, NIH3T3, HCT116, and MDA-MB-231 cells were cultured in DMEM/high-glucose (HyClone) supplemented with 10% fetal bovine serum, 100 units/ml penicillin and 100 μg/ml streptomycin (Sangon Biotech) at 37 °C in 5% CO2. MCF7 cells were cultured in MEM/high-glucose (HyClone), A549 cells were cultured in RPMI-1640 high-glucose medium (HyClone), and the other culture conditions were the same as those used for HCT116 cells. Corning TC-treated Culture Dish was used for routine cell culture. For the cell culture with 2D polyacrylamide-based hydrogels, hydrogels of high (40.40 ± 2.39 kPa) or low (1.00 ± 0.31 kPa) stiffness was generated according to the published protocol (Tse & Engler, 2010). Fibronectin solution (3μg/ml) and Sigmacote were used to coat the surface of the hydrogels. PEI (Polysciences) and Lipofectamine 2000 (Invitrogen) were used for plasmid transfection. Lipofectamine RNAiMAX (Invitrogen) was used for siRNA transfection.

### Plasmids and Reagents

Full-length PNRC1 cDNA was inserted into the pQCXIH vector, and the mutant pQCXIH-FLAG-PNRC1^W300A^ plasmid was constructed by using a KOD mutagenesis kit (Toyobo, Osaka, Japan) according to the manufacturer’s instructions. The pBABE-FLAG-YAP^5SA^/YAP^5SA-S94A^ and pRK7-FLAG-YAP^5SA^/YAP^5SA-S94A^ plasmids were obtained from laboratory storage. To generate the shRNA constructs targeting human TAZ, the targeting sequences were inserted into the pLKO.1-puro vector. The shRNA constructs targeting human YAP, SAMD4A, AJUBA, DDX6, DCP1A and LSM14A were generated by using the pLKO.1-hygro vector. The PNRC1 promoter and intron reporter plasmids and TEAD binding site mutant reporter plasmids were constructed by using the pGL3-Basic vector. siRNA/shRNA was used for all RNA silencing experiments in the study. The siRNA oligos targeting CHD4, RBBP4, PNRC1, SAMD4A, AJUBA, YAP, and TAZ were synthesized by Shanghai Genepharma Co., Ltd. The shRNA and siRNA targeting sequences and the primers used for plasmid construction are listed in Appendix Table S3. The following antibodies were used in this study: anti-FLAG (D6W5B, CST), anti-FLAG (M2, Sigma), anti-YAP/TAZ (D24E4, CST), anti-YAP (sc-101199, Santa Cruz), anti-TEAD4 (ab58310, Abcam), anti-CHD4 (14173-1-AP, Proteintech), anti-DDX6 (A9634, ABclonal), anti-DCP1A (A6824, ABclonal), anti-LSM14A (18336-1-AP, Proteintech), anti-AJUBA (A22039, ABclonal), anti-SAMD4A (17387-1-AP, Proteintech) and anti-PNRC1 (51052-1-AP, Proteintech).

### Cell proliferation, colony formation and cell migration assays

For the proliferation assay, cells (1×10^3^ per well) were seeded into a 96-well plate and cultured for 5 days. Cell viability was measured every day with a Cell Counting Kit 8 (CCK8) (Vazyme) according to the manufacturer’s instructions. Briefly, 10 μl of CCK8 reagent was added to each well and incubated for 2 hours. The absorbance at 450 nm was measured with a microplate reader to determine the relative numbers of viable cells. For the colony formation assay, cells (1×10^3^ per well) were seeded and cultured in six-well plates for 2 weeks. Then, the cells were stained with 1% crystal violet, and the number of colonies in each well was counted. For the cell migration assay, cells (1.5×10^5^ per well) in DMEM/high-glucose containing 0.1% FBS were seeded in the upper compartment of a Transwell chamber, while DMEM/high-glucose containing 10% FBS was placed in the lower compartment. After 60 hours, migrated cells were stained with 1% crystal violet and counted.

### Cell cycle and cell apoptosis assays

Cell cycle and cell apoptosis assays were performed according to the manufacturer’s instruction (FITC Annexin V Apoptosis Detection Kit I, BD, 556547; PI/RNase staining buffer, BD, 550825).

### qPCR, ChIP and luciferase reporter assays

qPCR and ChIP were performed as previously described (Zhu et al, 2020). For the luciferase assay, HEK293T cells were seeded in 24-well plates, incubated overnight to 50% confluence and then cotransfected with the PNRC1 luciferase reporter plasmid and the FLAG-YAP^5SA^ or FLAG-YAP^5SA-S94A^ plasmid. A Renilla luciferase plasmid was used as the control. After 24-36 hours, luciferase activity was measured by using a dual-luciferase reporter assay (Promega) and normalized to Renilla luciferase activity. For the luciferase assay using the YAP/TAZ knockdown of HCT116 cells, stable HCT116 cells were seeded in 24-well plates and cotransfected with the PNRC1 luciferase reporter and Renilla luciferase plasmids for 6h. Then, cells were replated to 6-well plates and cultured for 24- 36 hours at a low cell density before measuring the luciferase activity.

### Immunofluorescence staining

Cells were seeded in glass-bottom cell culture dishes one night before IF staining. The cells were washed with PBS and fixed with 4% paraformaldehyde. After the cells were permeabilized with 0.1% Triton X-100 at RT for 5 min, they were blocked with 3% BSA at RT for 1 hour. Then, the cells were incubated overnight with a rabbit anti- DDX6/DCP1A/LSM14A antibody (1:250) or a mouse anti-FLAG (M2) antibody (1:150) or mouse anti-YAP antibody (1:100). After a 1-hour incubation with Cy3-conjugated mouse and Alexa Fluor 488-conjugated rabbit secondary antibodies followed by a 1 min incubation with 0.1 μg/mL DAPI, the cells were visualized with an Olympus IX81 microscope.

### Xenograft assay and immunohistochemistry

Nude mice (4-6 weeks old, male) were obtained from SLAC Laboratory Animals LLC, Shanghai, China. All mouse procedures were approved by the Xinhua Hospital Animal Care and Use Committee. Male nude mice (4–6 weeks old) were randomly divided into four groups (n=6 mice per group) and injected in the right flank with 2 × 10^6^ of the indicated stable HCT116 cells resuspended in 100 μl of phosphate-buffered saline (PBS). Mice were sacrificed on Day 21, and the xenograft tumors were removed, photographed, and paraffin embedded for sectioning. The xenograft tumors were sectioned for H&E staining and immunohistochemical staining with anti-ki67 and anti-PNRC1 antibodies as described previously (Zhu et al, 2020).

### Colorectal cancer specimen

Patients with colorectal cancer who underwent curative surgery without prior treatments at the Department of Colorectal and Anal Surgery, Xinhua Hospital, Shanghai Jiao Tong University School of Medicine, between January 2008 and December 2018 were enrolled. Institutional review board approval and informed consent were obtained for all sample collections. None of the patients had any history of other tumors. Tumors and paired paracancerous normal tissues were collected during surgery. The generation of the CRC tissue array has been described and the IHC analysis of the CRC tissue array was performed according to our previous study (Zhu et al, 2020).

### Statistical analysis

Statistical analysis was performed by using GraphPad Prism 8.0.2 and SPSS 22.0. Typically, differences between two groups were evaluated using two-tailed Student’s *t* test or the chi-square test as indicated in the figure legends. One-way ANOVA was used for the experiments with more than two groups. Mann–Whitney U test and Kruskal-Wallis test were used for statistical analysis of the foci number in IF staining assay. Two-way ANOVA was used for statistical analysis of CCK8 assay. Paired Student’s t test was performed to assess the statistical significance of differential PNRC1 mRNA expression in 16 pairs of CRC and adjacent normal tissues. Pearson correlation analysis was used to statistically assess correlations of mRNA levels or codependencies from the DepMap database. The results are shown as averages; the error bars indicate the standard deviations (SDs). P values less than 0.05 were considered significant (*P < 0.05; **P < 0.01; ***P < 0.001; ****P < 0.0001).

## Supporting information

Appendix Table S1. The expression of P-body genes in YAPTAZ knockdown HCT116 cells

Appendix Table S2. Top 100 correlation for EDC4 in CRISPR (DepMap 22Q1 Public+Score, Chronos)

Appendix Table S3. siRNA, shRNA and primers used in this study

## Acknowledgments

This work was sponsored by the National Key R&D Program of China (2019YFC1316002), the National Natural Science Foundation of China (82172916, 82073056, 82273022), the Program for Professor of Special Appointment (Eastern Scholar) at Shanghai Institutions of Higher Learning (to C-Y. L.).

## Author contributions

C-Y.L., YL and WY conceived the project and prepared the manuscript. XS, XP and YGG performed most of the experiments and analyzed data. ZJD performed the bioinformatic analysis. LC performed the IHC analysis of the clinical tissues.

## Expanded View Figure legends

**Figure 1-Figure Supplement 1.**
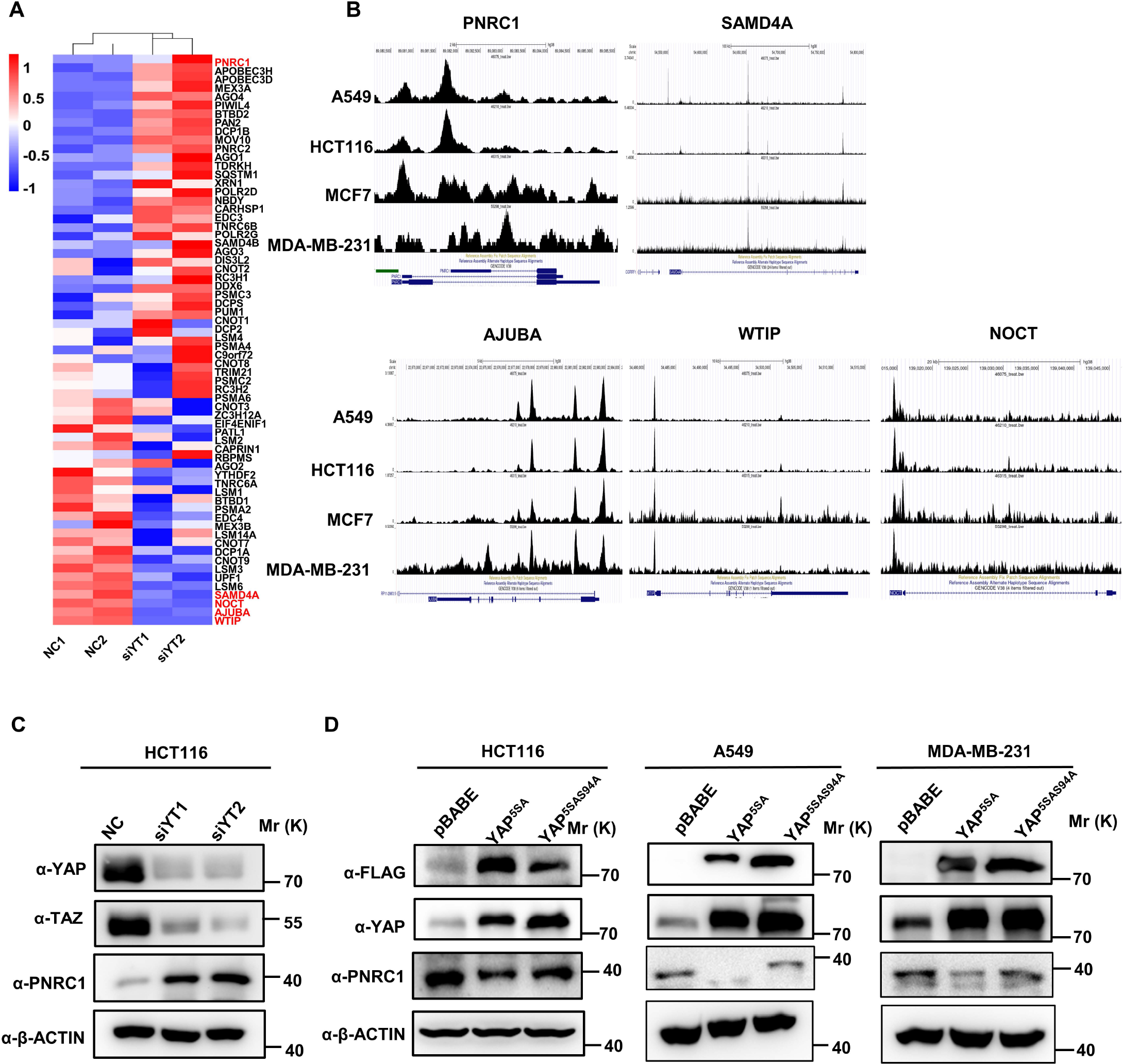
Supplemental figure related to Figure 1 (A) Heatmap showing the mRNA levels of 76 genes annotated as P-body-related genes, as detected in HCT116 cells by RNA-seq (GSE176475). (B) Representative sequencing TEAD4 ChIP-seq tracks at the SAMD4A/AJUBA/WTIP/NCOT/PNRC1 loci in HCT-116, A549, MDA-MB-231 and MCF7 cells. The ChIP-seq data were extracted from the Cistrome database and uploaded to the UCSC Genome Browser for visualization. (C) Western blot analysis of PNRC1, YAP, and TAZ in control and YAP/TAZ knockdown HCT116 cells. (D) Western blot analysis of PNRC1, YAP, and FLAG in control HCT116 cells and HCT116 cells overexpressing FLAG-YAP^5SA^ and YAP^5SA-S94A^. These data (C, D) are representative of 2 independent experiments.

**Figure 2-Figure Supplement 1.**
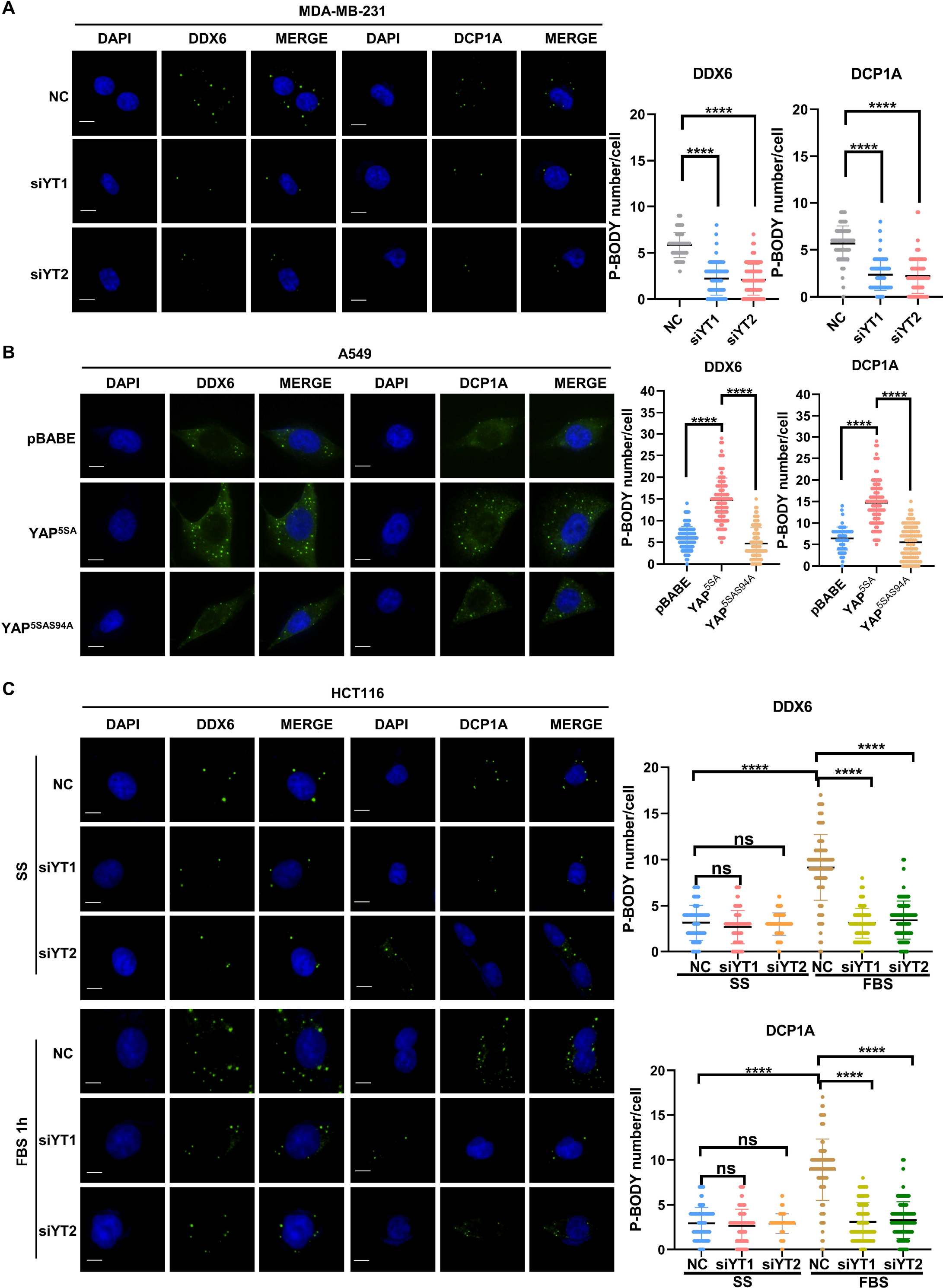
Supplemental figure related to Figure 2 (A) Immunofluorescence analysis of the P-body markers DDX6 and DCP1A in YAP/TAZ knockdown MDA-MB-231 cells. Cells were transfected with YAP/TAZ siRNA for 3 days before processing for immunofluorescence staining using anti-DDX6 and anti-DCP1A antibodies. Foci were counted in 100 cells per group. (B) Immunofluorescence analysis of DDX6 and DCP1A in A549 cells expressing YAP^5SA^ _and YAP_5SA-S94A_._ (C) Immunofluorescence analysis of DDX6 and DCP1A in HCT116 cells. Cells were transfected with control and YAP/TAZ siRNA for 3 days before overnight serum starvation. Then, the starved cells were treated with 10% FBS for 1 h before immunofluorescence staining. Kruskal-Wallis test was performed to assess statistical significance. These data (A-C) are representative of 3 independent experiments.

**Figure 2-Figure Supplement 2.**
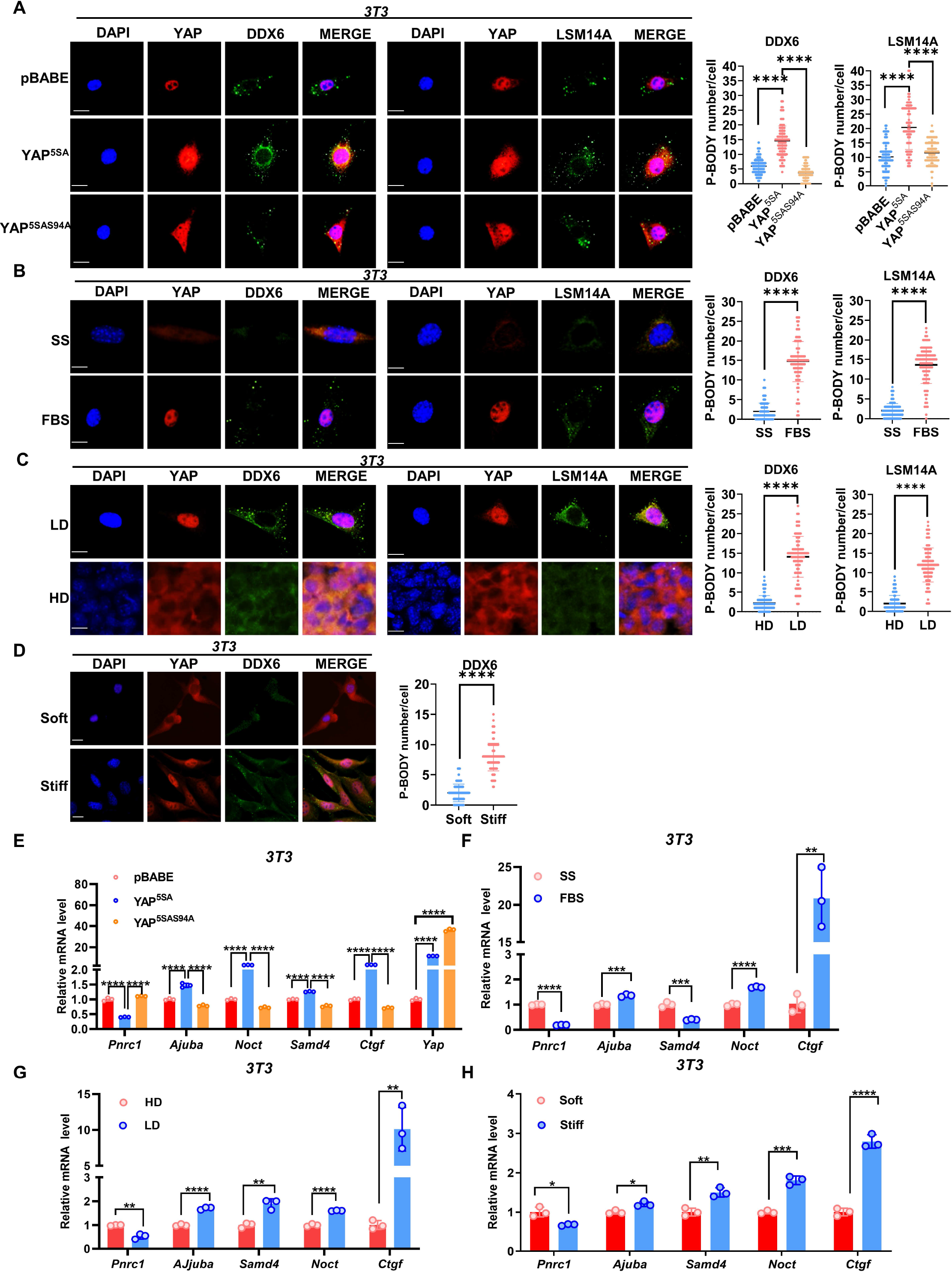
Supplemental figure related to Figure 2 (A) Immunofluorescence analysis of DDX6 and LSM14A in NIH3T3 cells expressing YAP^5SA^ and YAP^5SA-S94A^. Foci were counted in 100 cells per group. (B) Immunofluorescence analysis of DDX6 and LSM14A in NIH3T3 cells. Sparse Cells were treated with 10% FBS for 1 h after overnight serum starvation (SS). (C) Immunofluorescence analysis of DDX6 and LSM14A in NIH3T3 cells in sparse or confluent culture. (D) Immunofluorescence analysis of DDX6 and LSM14A in NIH3T3 cells cultured on soft (1kPa) to stiff (40kPa) matrices. (E-H) qPCR analysis of the indicated genes in NIH3T3 cells expressing YAP^5SA^ and YAP^5SA-S94A^ (E) or NIH3T3 cells treated with 10% FBS for 1 h after overnight serum starvation (F) or cultured under sparse or confluent conditions in standard culture medium (G) or cultured on soft (1kPa) to stiff (40kPa) matrices (H). Kruskal-Wallis test (A), Mann– Whitney U test (B-D), one-way ANOVA (E) and Two-tailed Student’s t test (F-H) were performed to assess statistical significance. These data (A-H) are representative of 3 independent experiments.

**Figure 3-Figure Supplement 1.**
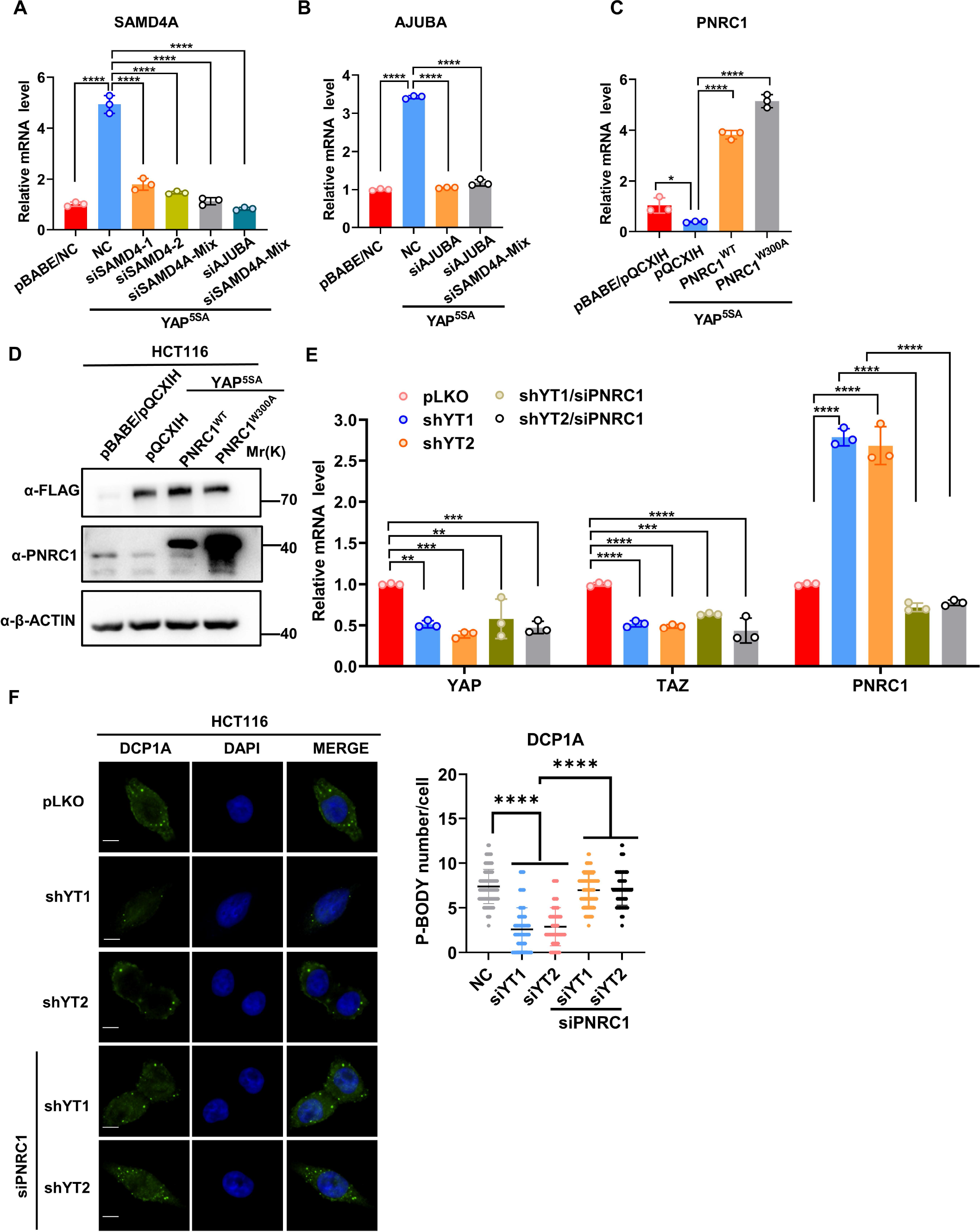
Supplemental figure related to Figure 3 (A, B) qPCR analysis of the mRNA levels of SAMD4A (A) and AJUBA (B) in HCT116 cells stably expressing FLAG-YAP^5SA^ and YAP^5SA^ and transfected with SMAD4A or AJUBA siRNA. n=3 biologically independent samples per group. (C, D) qPCR (C) and Western blot (D) analysis of PNRC1 expression in HCT116 cells stably expressing FLAG-YAP^5SA^ alone or in combination with PNRC1^WT^ or PNRC1^W300A^. (E) qPCR analysis of the mRNA level of PNRC1/YAP/TAZ in YAP/TAZ knockdown HCT116 cells transfected with PNRC1 siRNA. n=3 biologically independent samples per group. (F) Immunofluorescence analysis of DCP1A in YAP/TAZ knockdown HCT116 cells transfected with PNRC1 siRNA. Foci were counted in 100 cells per group. One-way ANOVA (A-C, E) and Kruskal-Wallis test (F) were performed to assess statistical significance for qPCR analysis in this figure. The data (F) is representative of 2 independent experiments.

**Figure 4-Figure Supplement 1.**
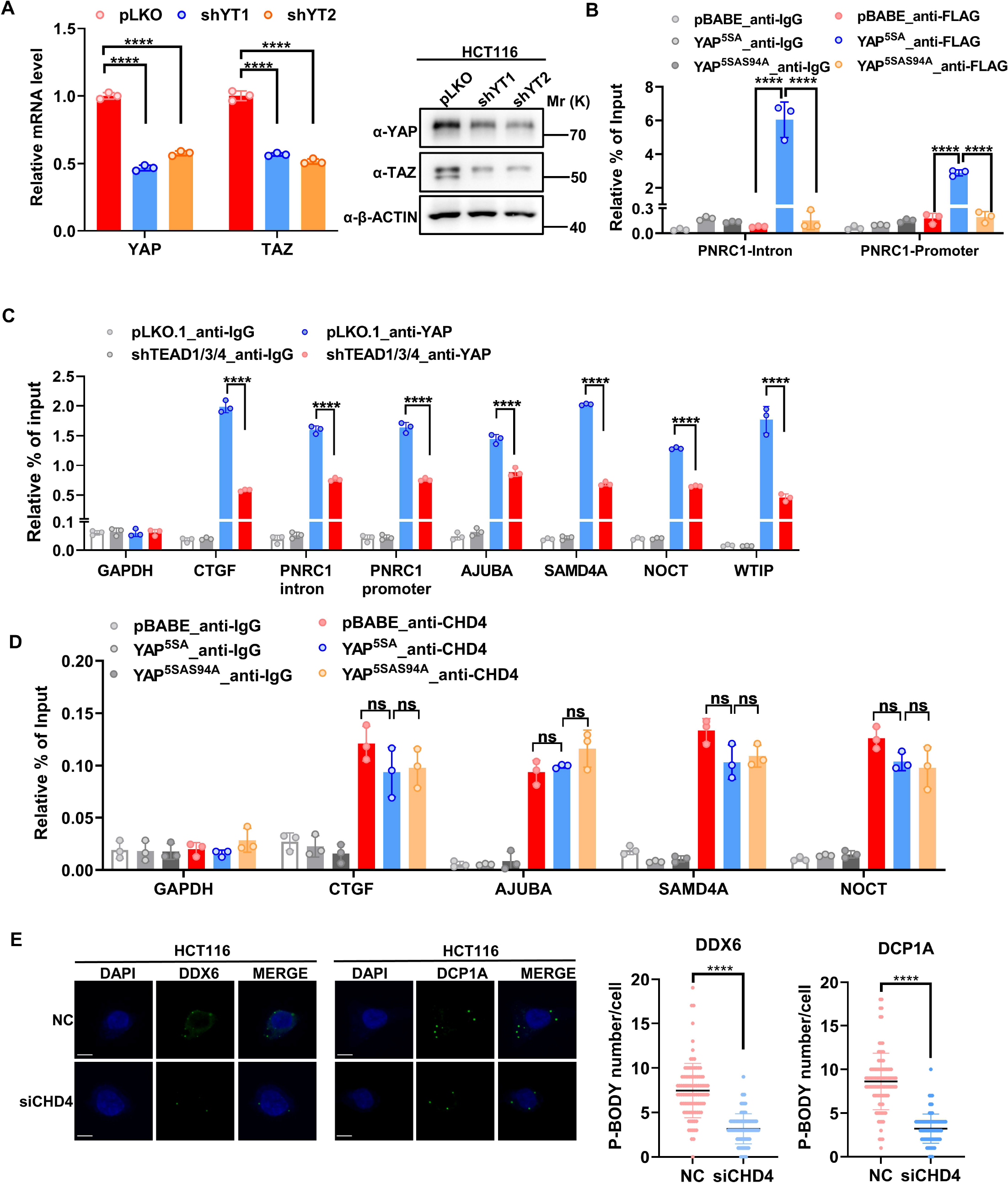
Supplemental figure related to Figure 4 (A) qPCR and WB analysis of the YAP/TAZ knockdown stable HCT116 cells. (B) ChIP–qPCR analysis of FLAG-YAP^5SA^, YAP^5SA-S94A^ binding to the PNRC1 promoter and intronic regions in control and HCT116 cells stably expressing FLAG-YAP^5SA^ or _YAP_5SA-S94A_._ (C) ChIP–qPCR analysis of YAP binding to the genomic locus of the indicated P-body-related genes in control and HCT116 cells with stably knockdown of TEAD1/3/4. (D) ChIP–qPCR analysis of CHD4 binding to the YAP binding sites of indicated genes in control and HCT116 cells stably expressing FLAG-YAP^5SA^ or YAP^5SA-S94A^. One-way ANOVA was performed to assess statistical significance for qPCR analysis in this figure. These data (B-D) are representative of 2 independent experiments. (E) Immunofluorescence analysis of DDX6 and DCP1A in HCT116 cells. Cells were transfected with control and CHD4 siRNA for 3 days before Immunofluorescence analysis. Foci were counted in 100 cells per group. Kruskal-Wallis test was performed to assess statistical significance.

**Figure 5-Figure Supplement 1.**
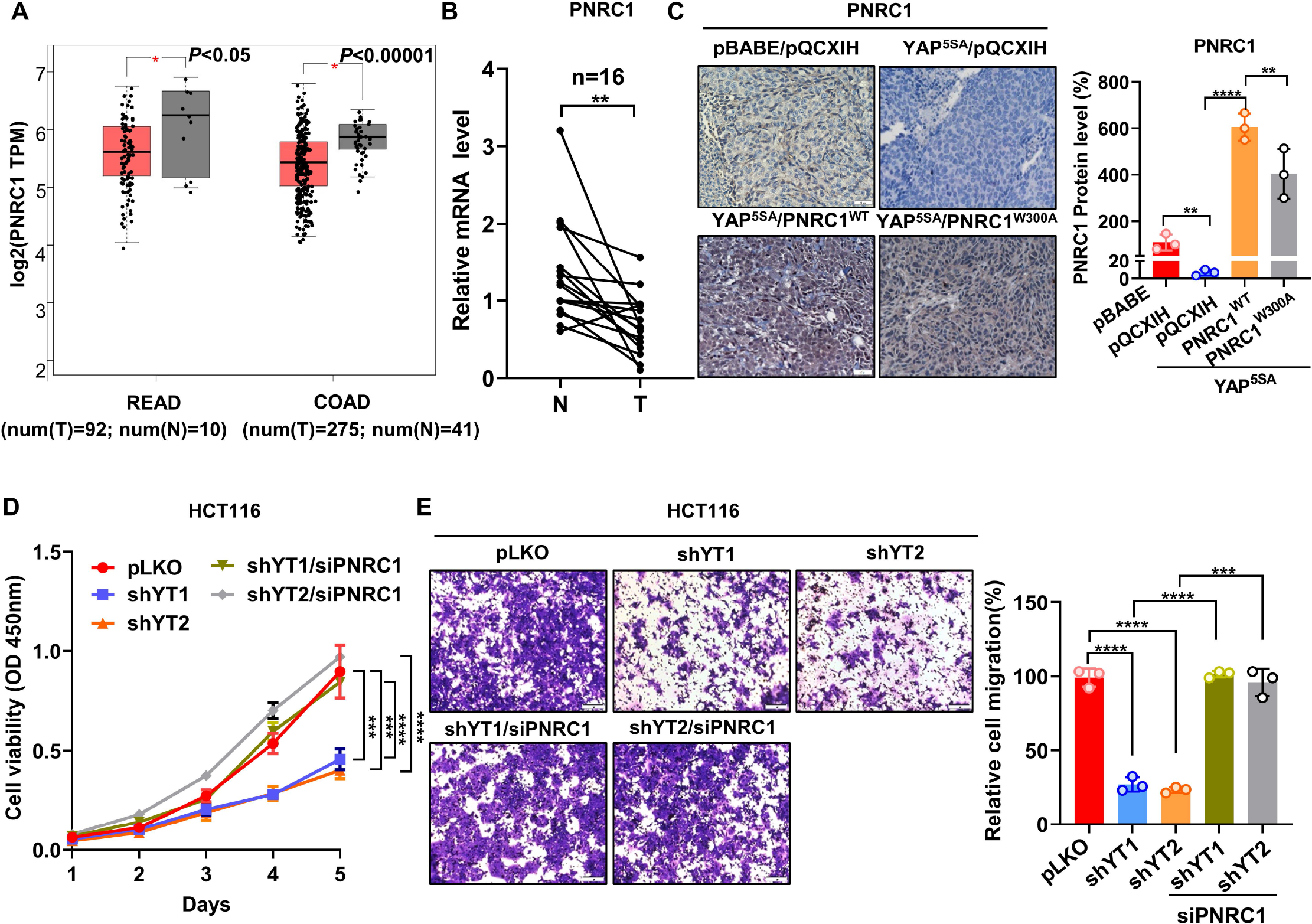
Supplemental figure related to Figure 5 (A) Downregulated mRNA expression of PNRC1 in colon cancer (COAD) and rectal cancer (READ) CRC datasets in TCGA. The mRNA levels of PNRC1 were extracted from the GEPIA database. (B) qPCR analysis of the mRNA levels of PNRC1 in 16 paired normal mucosa and colorectal tumor tissues. Paired Student’s t test was performed to assess statistical significance. (C) Representative images of IHC staining of PNRC1 in xenograft tumors formed from HCT116 cells stably expressing YAP^5SA^ alone or in combination with PNRC1^WT^ or PNRC1^W300A^ (n=3). (D, E) CCK8 proliferation assay (n=4) (D) and Transwell migration assay (n=3) (E) of YAP/TAZ knockdown HCT116 cells transfected with PNRC1 siRNA. One-way ANOVA (A, C, E) and two-way ANOVA (D) were performed to assess statistical significance. These data (D, E) are representative of 3 independent experiments.

**Figure 6-Figure Supplement 1.**
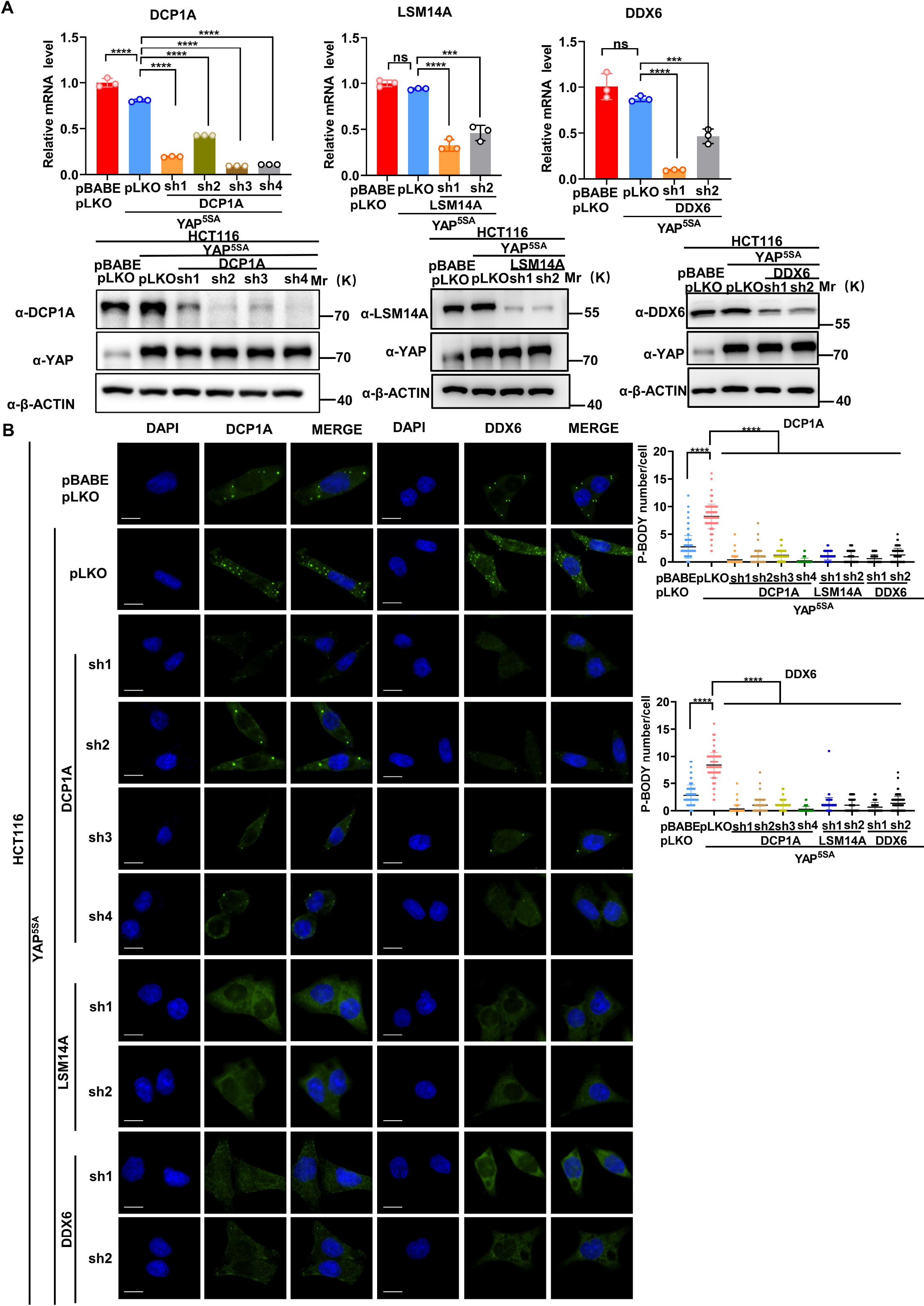
Supplemental figure related to Figure 6 (A) qPCR and WB analysis of the knockdown efficiency of DCP1A, LSM14A and DDX6 in HCT116 cells stably expressing YAP^5SA^ with or without knockdown of DCP1A, LSM14A or DDX6. n=3 biologically independent samples per group. (B) Immunofluorescence analysis of DDX6 and DCP1A in HCT116 cells stably expressing YAP^5SA^ with or without knockdown of DCP1A, LSM14A or DDX6. Foci were counted in 100 cells per group. One-way ANOVA (A) and Kruskal-Wallis test (B) were performed to assess statistical significance.

**Figure 6-Figure Supplement 2.**
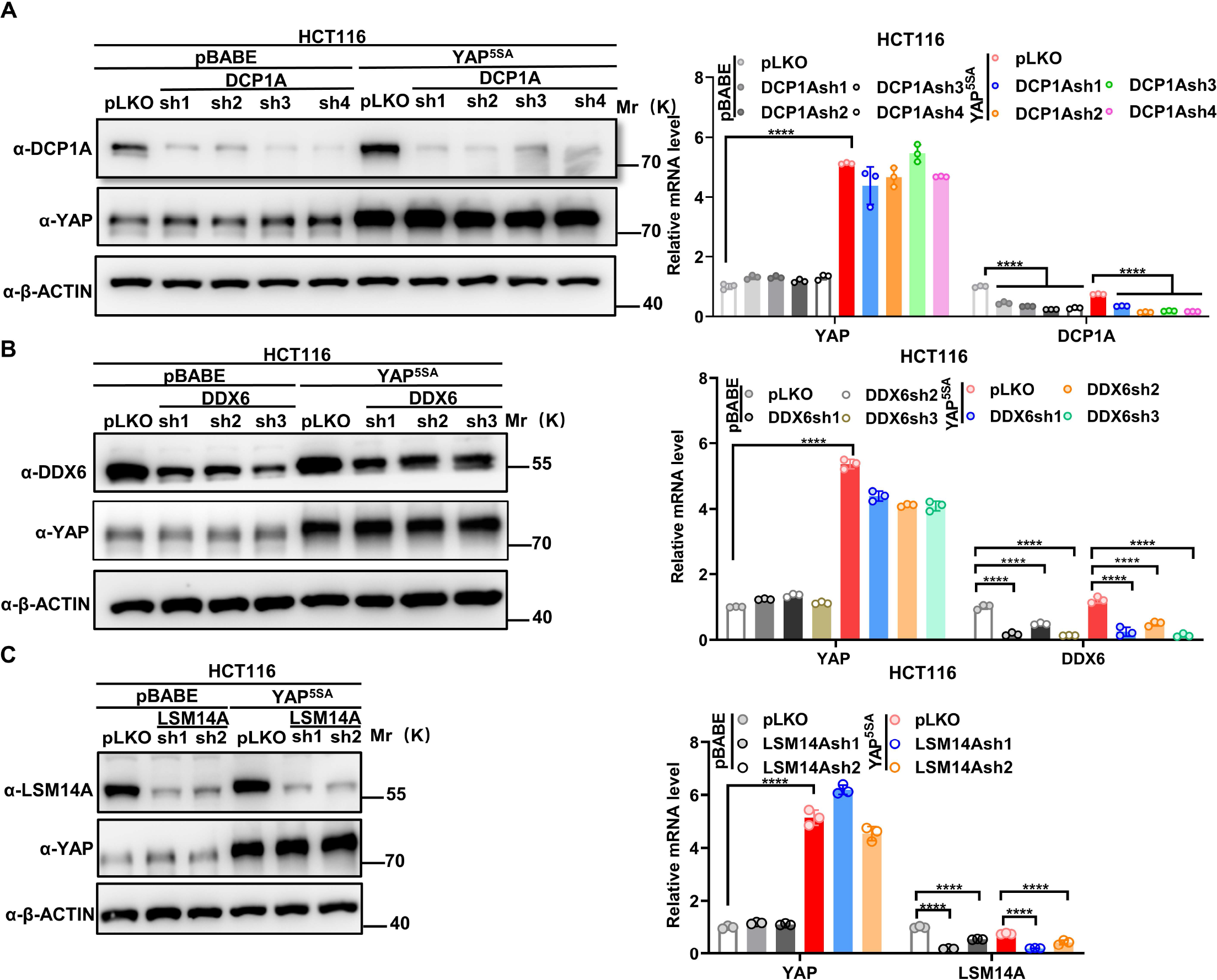
Supplemental figure related to Figure 6 (A-C) qPCR and WB analysis of the knockdown efficiency of DCP1A (A), DDX6 (B) and LSM14A (C) in control HCT116 cells with or without knockdown of DCP1A, LSM14A or DDX6 and stable YAP^5SA^ overexpression HCT116 cells with or without knockdown of DCP1A, LSM14A or DDX6. n=3 biologically independent samples per group. One-way ANOVA was performed to assess statistical significance.

**Figure 6-Figure Supplement 3.**
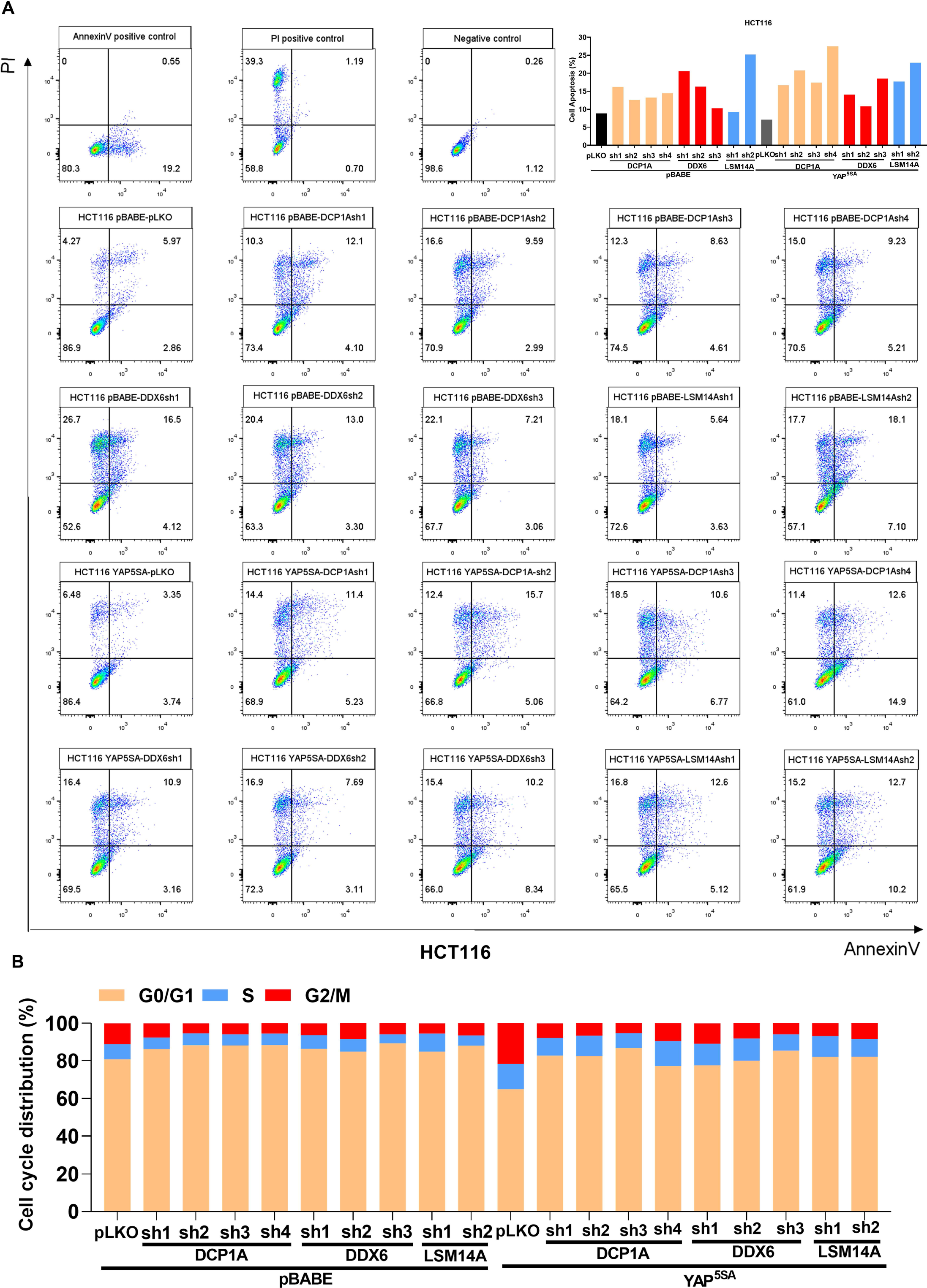
Supplemental figure related to Figure 6 (A, B) Cell apoptosis (A) and cell cycle (B) analysis of the control HCT116 cells with or without knockdown of DCP1A, LSM14A or DDX6 and stable YAP^5SA^ overexpression HCT116 cells with or without knockdown of DCP1A, LSM14A or DDX6.

**Figure 6-Figure Supplement 4.**
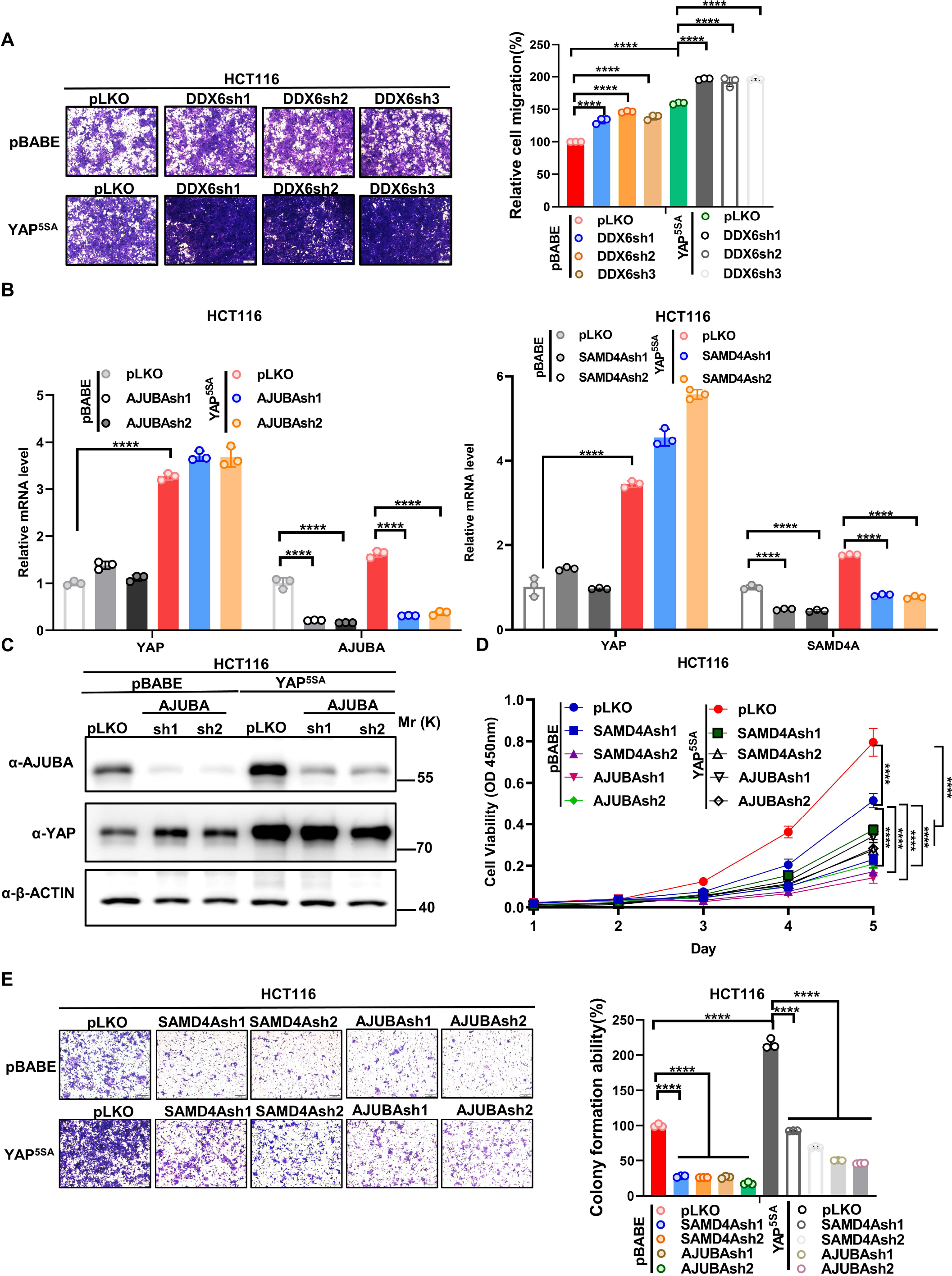
Supplemental figure related to Figure 6 (A) Transwell assay of control HCT116 cells with or without knockdown of DDX6 and HCT116 cells stably expressing YAP^5SA^ with or without knockdown of DDX6. n=3 biologically independent samples per group. (B, C) qPCR (B) and WB (C) analysis of the knockdown efficiency of AJUBA and SAMD4A in control HCT116 cells with or without knockdown of AJUBA or SAMD4A and stable YAP^5SA^ overexpression HCT116 cells with or without knockdown of AJUBA or SAMD4A. n=3 biologically independent samples per group. (D, E) CCK8 proliferation (n=5) (D) and transwell (E) assays of control HCT116 cells with or without knockdown of AJUBA or SAMD4A and stable YAP^5SA^ overexpression HCT116 cells with or without knockdown of AJUBA or SAMD4A. one-way ANOVA (A, B, E) and Two-way ANOVA (D) were performed to assess statistical significance in this figure. These data (A, D, E) are representative of 3 independent experiments.

**Figure 7-Figure Supplement 1.**
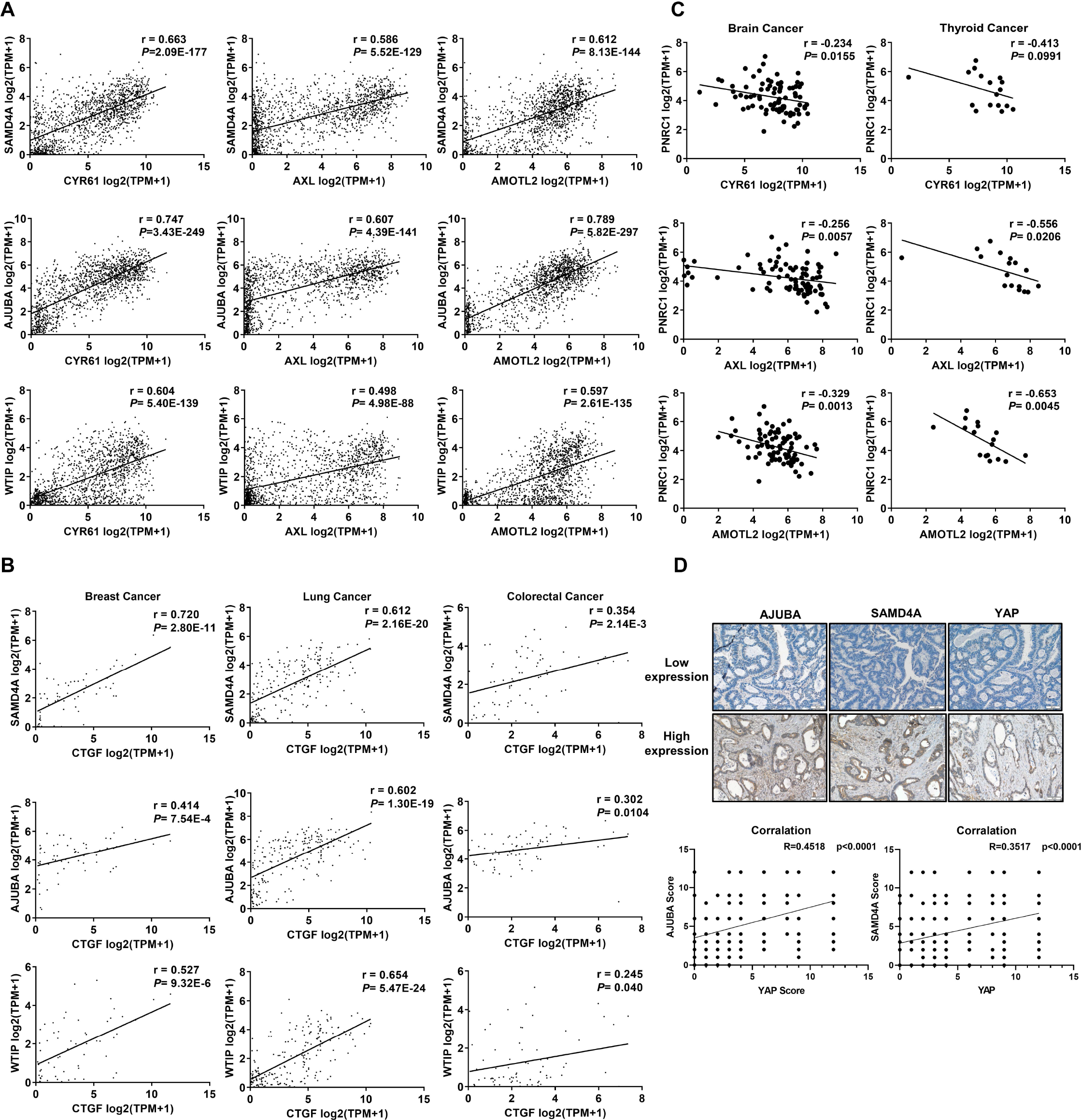
Supplemental figure related to Figure 7 (A) Positive correlations between the mRNA levels of YAP target genes (CYR61, AXL, and AMOTL2) and SAMD4A/AJUBA/WTIP in 1393 cancer cell lines. (B) Positive correlations between the mRNA levels of YAP target genes (CTGF) and SAMD4A/AJUBA/WTIP in brain cancer (n=83), lung cancer (n=207) and colorectal cancer (n=71) cell lines. (C) Negative correlations between the mRNA levels of YAP target genes (CYR61, AXL, and AMOTL2) and PNRC1 in brain cancer cell lines (n=83) and thyroid cancer cell lines (n=17). (D) IHC analysis of AJUBA, SAMD4A and YAP protein expression in colorectal cancer tissues (n= 294). Pearson (A-C) and spearman (D) correlation analysis was used to assess statistical significance.

**Figure 7-Figure Supplement 2.**
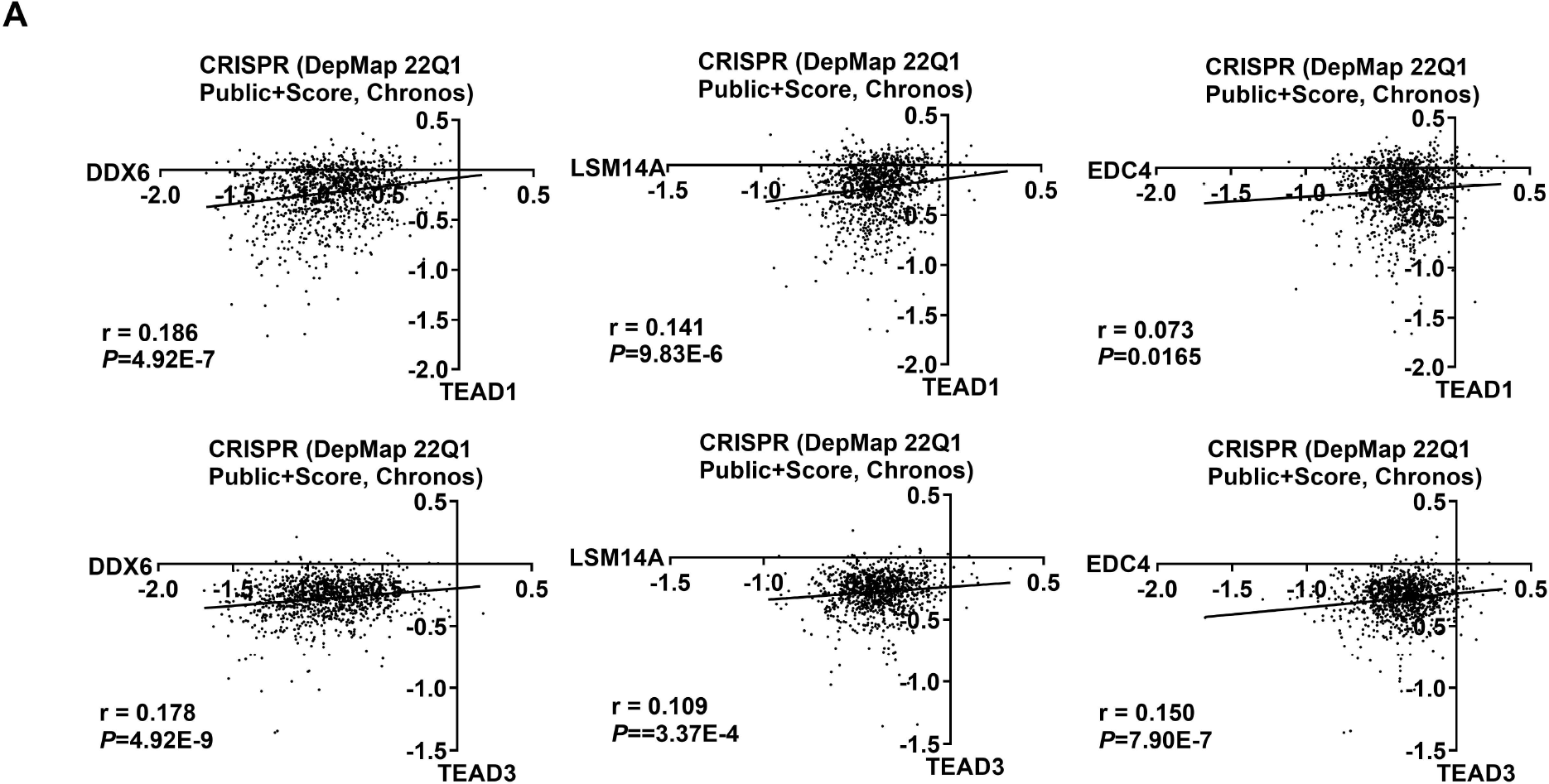
Supplemental figure related to Figure 7 (A) Positive correlations between the dependency scores of TEAD1/3 and DDX6/LSM14A/EDC4 in 1070 cancer cell lines. The Chronos dependency scores were extracted from the DepMap database. The negative Chronos score indicates decreased cell proliferation upon gene knockout. Pearson correlation analysis was used to assess statistical significance.

**Figure 7-Figure Supplement 3-5.**
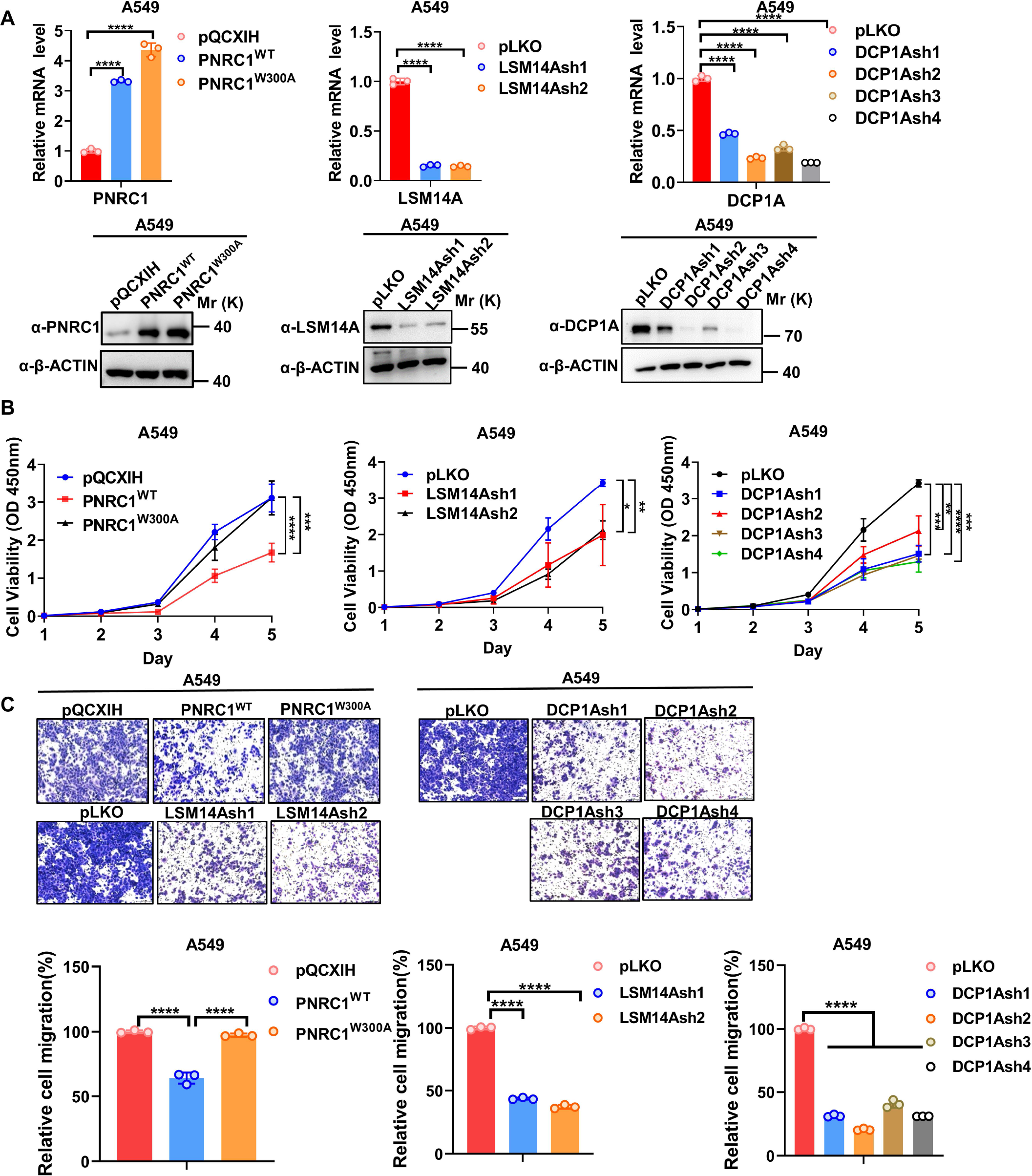
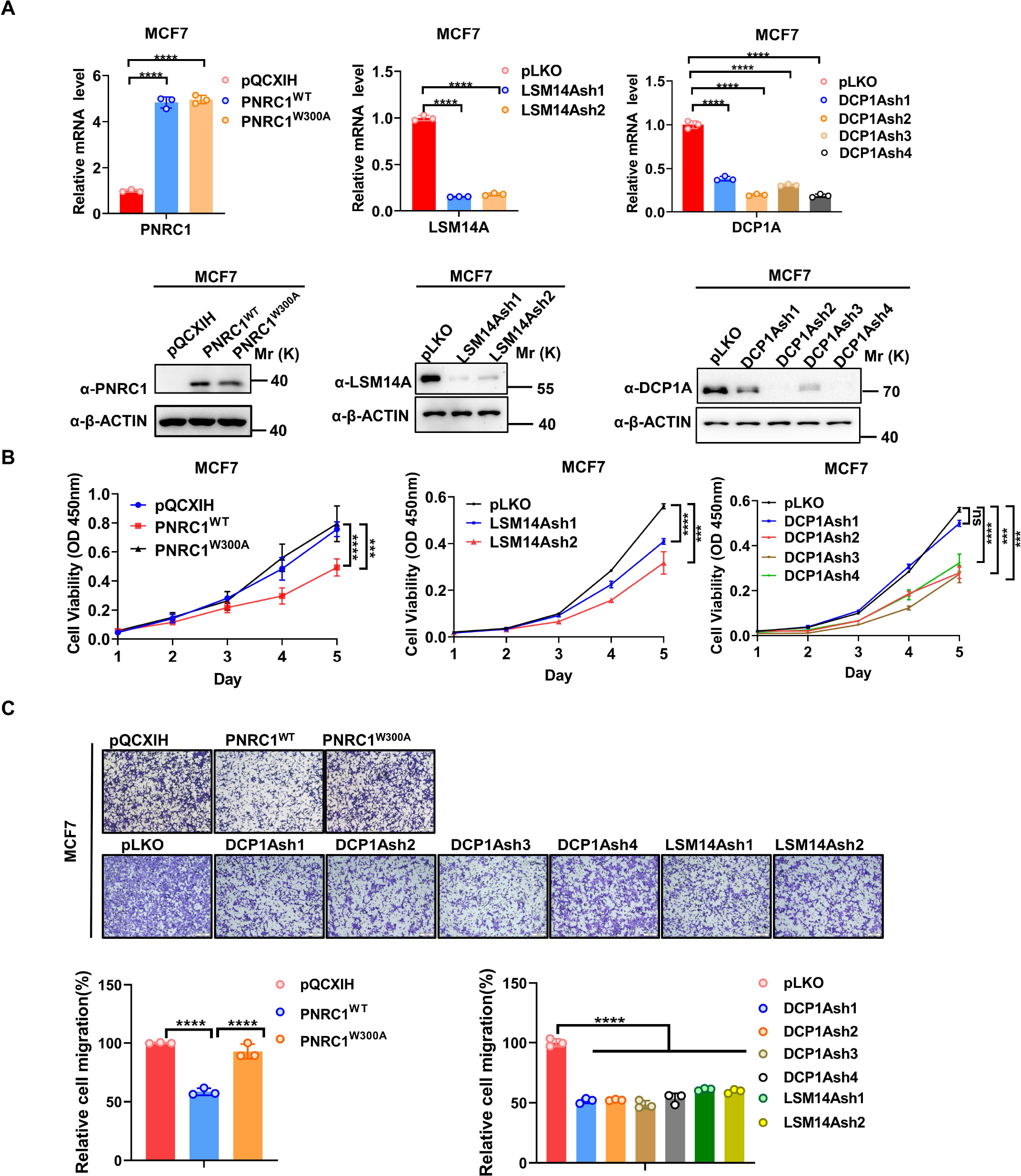
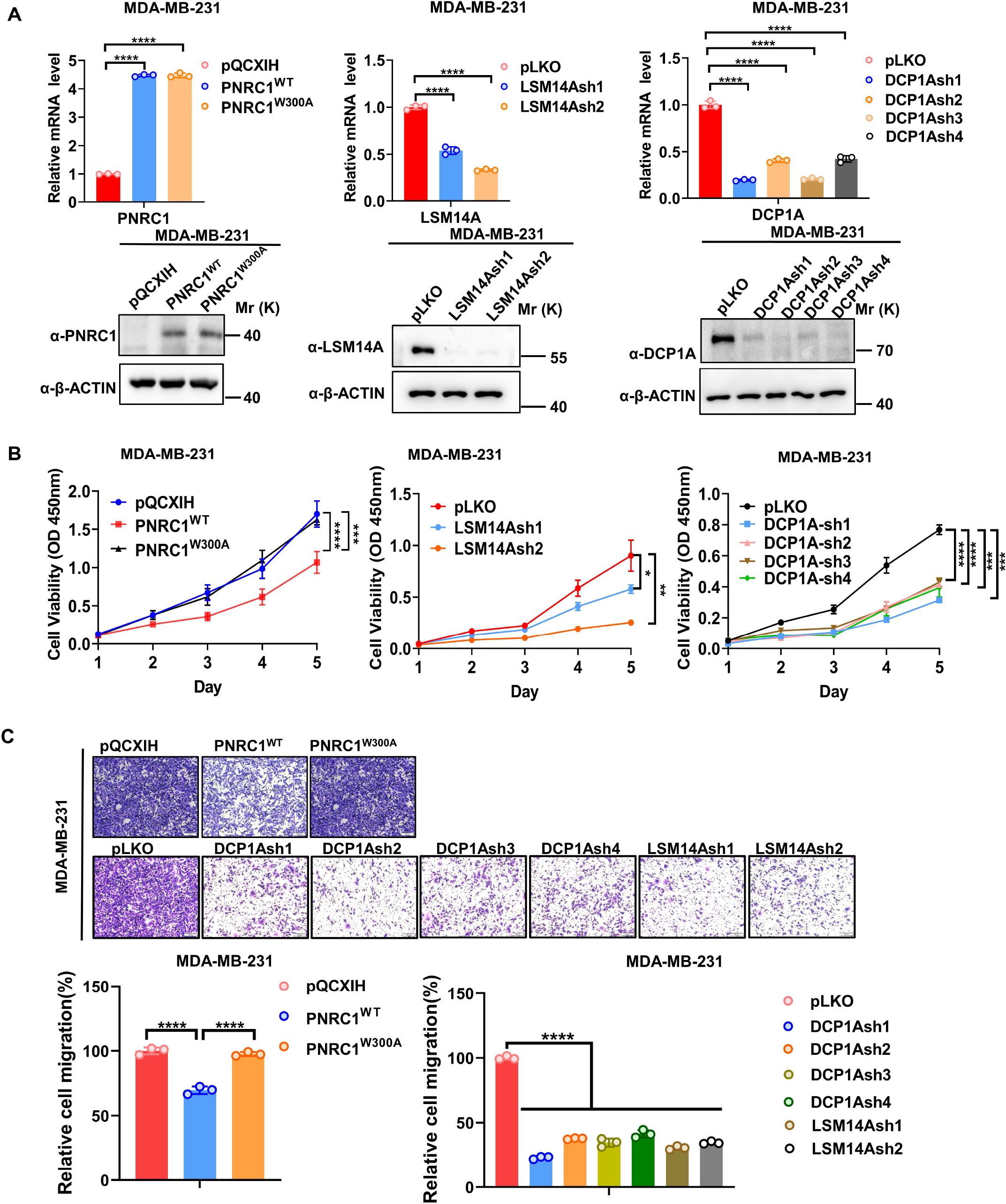
Supplemental figure related to Figure 7 (A) qPCR and WB analysis of the knockdown efficiency of DCP1A / LSM14A or PNRC overexpression in A549 (Figure 7-Figure Supplement 3), MCF7 (Figure 7-Figure Supplement 4) and MDA-MB-231 (Figure 7-Figure Supplement 5) cells stably expressing PNRC1^WT^/PNRC1^W300A^ or with knockdown of DCP1A and LSM14A. (B, C) CCK8 proliferation (n=5 for PNRC1 OE, n=3 for knockdown of DCP1A and LSM14A) (B) and transwell (C) assays of A549 (Figure 7-Figure Supplement 3), MCF7 (Figure 7- Figure Supplement 4) and MDA-MB-231 (Figure 7-Figure Supplement 5) cells stably expressing PNRC1^WT^/PNRC1^W300A^ or with knockdown of DCP1A and LSM14A. Two-way ANOVA (B) and one-way ANOVA (C) were performed to assess statistical significance in this figure. These data (B, C) are representative of 2 independent experiments.

